# Actin polymerization and crosslinking drive left-right asymmetry in single cell and cell collectives

**DOI:** 10.1101/2021.04.22.440942

**Authors:** Y. H. Tee, W. J. Goh, X. Yong, H. T. Ong, J. Hu, I. Y. Y. Tay, S. Shi, S. Jalal, S. F. H. Barnett, P. Kanchanawong, W. Huang, J. Yan, V. Thiagarajan, A. D. Bershadsky

## Abstract

Deviations from mirror symmetry in the development of bilateral organisms are highly stereotypic and genetically predetermined, but their mechanisms are not sufficiently understood. At the cellular level, self-organization of the actin cytoskeleton results in chiral actin swirling, and cells in groups confined to micropatterns demonstrate chiral cell alignment. The relationship between individual and collective cell chirality is unclear, and molecular players involved remain essentially unidentified. Here, by screening major actin-associated proteins and deep-learning-based morphometric analysis of actin patterns, we found that knockdowns of specific actin polymerization regulators, such as mDia1, ARPC2, and cofilins 1&2, abolished chiral actin swirling, while depletion of profilin 1 and CapZβ, reversed its direction in an actin crosslinker α-actinin1-dependent manner. Analysis of these and other knockdowns and pharmacological treatments revealed a robust correlation between their effects on the chirality of individual cells and confined cell groups. Thus, actin-driven cell chirality may underlie tissue and organ asymmetry.

**One Sentence Summary:** Cell chirality determined by specific regulators of actin polymerization drives left-right asymmetry emergence in cell groups

## Introduction

While the majority of multicellular organisms demonstrate approximate bilateral symmetry, many important features of the body layout such as position of visceral organs, as well as the shape of organs themselves are usually asymmetric. These asymmetries are tightly programmed and aberrations in such a program can lead to severe defects in embryonic development (*1*). Several examples of left-right asymmetry emerging have been discovered. At the level of entire organisms, emergence of left-right asymmetry has been thoroughly described for visceral organs in vertebrates (*1, 2*) and formation of asymmetric body in snails (*3*). The processes of asymmetric organogenesis include heart-looping in vertebrates (*4*), chiral shaping of hindgut and genitalia (*5, 6*), and asymmetric tilting of wing bristles in *Drosophila* (*7*). Cell groups confined to micropatterned adhesive substrates (in the form of stripes or rings) demonstrated chiral cell alignment and movement (*8–10*). Finally, on the single cell level, the processes of intracellular swirling, cortical flow, and cell migration can demonstrate left-right asymmetry (*11–14*).

It is commonly believed that mechanisms underlying emergence of left-right asymmetry in diverse biological systems are based on the function of special chiral molecules (*15*), and in particular the chiral cytoskeletal fibers (*16*). Indeed, several classes of cytoskeletal proteins were shown to be involved in the processes of left-right asymmetry development listed above. While the asymmetric positioning of visceral organs depends on numerous cilia-related proteins (*17*) and attributed to cilia function in specialized cells located in the embryonic node (left-right organizer), actin cytoskeleton related proteins are involved in other examples of asymmetry. In particular, non-conventional myosins 1d and 1c are needed for asymmetric hindgut and male genitalia development in *Drosophila* (*5, 6*), and myosin 1d is sufficient to induce chirality in other *Drosophila* organs (*18*). We previously proposed the role of formin family proteins in development of actin cytoskeleton chirality (*11*) and several recent publications established the role of diaphanous formin in dextral snail chirality (*19, 20*), Daam formin in *Drosophila* hindgut and genitalia chirality (*21*), and *Caenorhabditis elegans* CYK-1 formin in chiral cortical flow of *C.elegans* zygote (*22*). Finally, some data indicated the involvement of the actin filament crosslinking protein alpha-actinin1 in emergence of chirality in individual cells and cell spheroids *in vitro (11, 23)*. Importantly, the helical nature of actin filaments allows all these proteins to interact with filaments in a chiral manner and produce different kinds of chiral movement and/or chiral superstructures (*18, 24, 25*). Nevertheless, how intrinsic chirality of the actin filaments is translated *in vivo* into chirality of cells, and whether this is sufficient to explain the emerging of chirality in multicellular groups such as tissue and organs remains obscure.

In particular, it is unknown whether left-right asymmetry emerges as a result of the activity of a single “chiral determinant” such as myosin 1d (*18*) (perhaps different proteins in different systems) or is mediated by coordinated activities of a group of proteins with complementary functions. Here, we addressed this question by systematic investigation of the involvement of major actin-associated proteins in the regulation of left-right asymmetry of the actin cytoskeleton in individual cells. In the course of this analysis, we examined the effects of depletion of chirality regulators found in previous studies, namely formins, myosin 1c and 1d, and alpha-actinin1. We show that even in this simple system the chiral swirling of actin depends on the functions of several groups of actin regulators. We reveal several types of such regulators: depletion of some of them reduced or abolished chirality, while depletion of others reversed the direction of chirality.

The discovery of numerous regulators of chiral morphogenesis in individual cells allowed us to perform detailed comparison between factors affecting individual and collective cell chirality. It was previously unclear whether chiral asymmetry of the actin cytoskeleton in individual cells is related to emergence of collective chirality in cell groups. Here, we found that the majority of treatments affecting the asymmetric self-organization of the actin cytoskeleton in individual cells also affected the asymmetric alignment of cell groups. In particular, all factors that reversed direction of cytoskeletal chirality in individual cells also reversed the direction of chiral cell alignment in cell groups. Altogether our findings provide the background for the future understanding of the processes of emerging left-right asymmetry in tissues and organs.

## Results

### Assessment of chiral organization of radial fibers

We have previously shown that human fibroblasts (HFF) plated on fibronectin-coated circular islands with an area of 1800 μm^2^ formed a chiral pattern of organization of actin filament bundles. Radial actin fibers originating from focal adhesions at the cell edge eventually tilted to the right from the axis connecting the focal adhesion with the cell center. This produced chiral anti clockwise swirling in the centripetal movement of transverse fibers along the tilted radial actin fibers (*11*) (Movie S1). To quantitatively investigate the molecular requirements for actin cytoskeleton chiral self-organization, we introduced a quantitative method of assessment of radial fiber tilting (Fig. 1, A to H). First, we segmented the radial fibers using a deep-learning network (Unet-ResNet50) (Fig. 1, A, B, E and F). Second, the image of cells on the pattern was subdivided into 8 concentric annular rings located at given distances from the cell edge (Fig. 1, C and G). The average tilt of radial fiber segments located in each ring was calculated (Fig. 1, D and H and S1). The radial fiber pattern was then characterized by the curve showing the average tilt of radial fiber segments as a function of their distance from the cell edge (Fig. 1L). In addition, to further characterize the variability between cells, we compared histograms characterizing the distribution of the average radial fiber tilt in the single annulus located between 6 – 10 or 8 – 12 microns from the cell edge (Fig. 1, I to K). Details of our methodology are described in methods section.

**Fig. 1.**
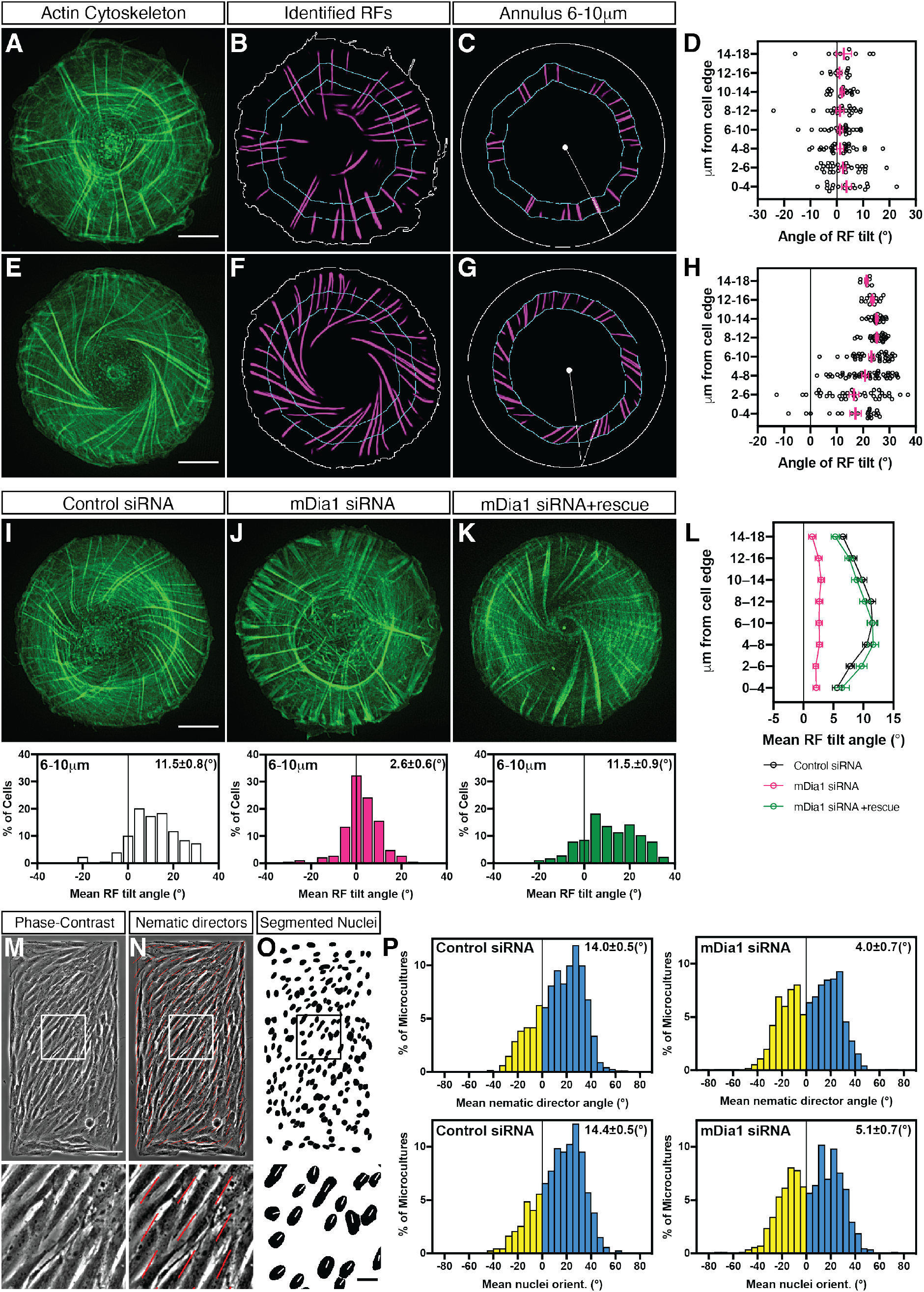
Quantification of left-right asymmetry in actin organization in individual cells and chiral alignment of cells in confined cell groups for control and mDia1 knockdown cells. (**A** to **H**) Measurements of the radial fiber (RF) tilt. Fluorescence images of actin cytoskeleton in radial (**A**) and chiral (**E**) cells obtained by phalloidin staining and RFs were identified by deep-learning procedure (**B** and **F**). The tilts of all RF segments in the concentric belts (annuli) located at given distance from the cell edge were measured as shown in (**C**) and (**G**). See also Fig. S1. The tilt values in all eight annuli corresponding to cell (**A**) and (**B**) are shown in (**D**) and (**H**) respectively. (**I** to **L**) Effect of mDia1 knockdown and rescue on RF tilt. Typical examples of actin organization in control siRNA (**I**), mDia1 siRNA (**J**) and mDia1 siRNA plus mDia1 full-length plasmid (**K**) transfected cells 6 hours following cell plating on circular patterns are shown. The histograms (**I** to **K**) show the distribution of average RF tilt in the 6-10 µm annulus in cells under corresponding conditions. (**L**) Average values of RF tilts (mean±*s.e.m*) as a function of the distance from the cell edge. mean±*s.e.m* values (in **I** to **L**) of these average RF tilts were obtained from 179 control cells, 186 mDia1 knockdown cells, and 177 mDia1 knockdown cells rescued by mDia1 plasmid overexpression. (**M** to **P**) Effect of mDia1 knockdown on left-right asymmetry of cell alignment in microcultures confined to rectangular micropatterns. (**M**) Phase-contrast image of cells 48 hours following plating on rectangular adhesive pattern (300×600μm). (**N**) Image shown in (**M**) overlayed with red lines representing average local orientation of cells (nematic directors). (**O**) Segmented Hoechst 33342-stained nuclei of cells shown in (**M**). Boxed areas in **M** to **O** are shown at higher magnification below. (**P**) Histograms showing distributions of the values of mean nematic directors (upper row) and mean nuclei orientation (lower row) characterizing individual microcultures. The histograms were built based on average nematic director angles from 1530 control and 894 mDia1 knockdown microcultures, or average nuclei orientation angles from 1521 control and 844 mDia1 knockdown microcultures respectively. Mean±*s.e.m* values are indicated at the top right corner of each histogram. Scale bars, 10μm (**A**, **E**, **I** to **K**); 100μm (**M**); 20μm (**O**, boxed region).

### Actin polymerases mDia1 and Arp2/3, and actin depolymerizing proteins cofilins 1 and 2 are required for development of a chiral actin pattern

We first systematically evaluated the effects of the knockdowns of 10 formin family members highly-expressed in fibroblasts (Fig. S2A) on chiral organization of radial fibers. Knockdown of diaphanous-related formin, mDia1, did not prevent the formation of either radial or transverse fibers, but abolished the tilting of the radial fibers (Fig. 1J). As a result, mDia1 depleted cells exhibited a radially symmetric organization of the actin cytoskeleton (Fig. 1, J and L). The inhibitory effect of mDia1 knockdown on chiral actin pattern formation can be rescued by expression of exogenous mDia1-GFP construct (Fig. 1, K and L and S2D). The results of knockdown of several other abundant formins can be classified into two groups (Fig. S2, B, C, E and F). FMNL2, FHOD3, Daam1 and FMN2 knockdowns reduced the degree of actin cytoskeleton chirality, albeit not to such extent as knockdown of mDia1 (Fig. S2B). The knockdowns of other examined formins (mDia2, mDia3, INF2, FHOD1 and Daam2) did not have an apparent effect on chirality (Fig. S2C).

We further studied the cell orientation in multicellular microcultures on rectangular adhesive islands with a 1:2 aspect ratio (300×600 μm). At 48 hours following plating, the cells approached confluency and aligned mainly along the diagonal of the rectangles as seen from the orientation of nuclei or local average cell orientation characterized by “nematic directors” on phase contrast images (*26*) (Fig. 1, M to O and Movie S2). Thus, each rectangular microculture was characterized by the mean angle between local nematic directors and the long axis of the rectangle (Fig. 1, N and P). In complementary set of measurements, the long axes of elliptic nuclei were used instead of nematic directors (Fig. 1, O and P). In control microcultures, the cells orientation was chiral so that the distribution of the values of angles characterizing individual rectangles was asymmetric (Fig. 1P). In other words, the cells on the rectangular pattern preferentially aligned into a “И”-orientation rather than a “N”-orientation (when observed from above). We also examined rectangles with other aspect ratios and found that the 300×600 μm size was optimal for asymmetric alignment of microcultures (Fig. S3, A to D).

We found that knockdown of mDia1 abolished the chirality of microculture orientation on rectangles so that distribution of the angles characterizing individual microcultures became bimodal (Fig. 1P), meaning that mDia1 knockdown cells in the rectangular microcultures oriented themselves in “И” or “N” fashion with equal probability. Examining the knockdowns of other formins revealed that FMNL2, FHOD3, Daam1, mDia2, and mDia3 knockdown slightly reduced the mean angles of both nematic directors and nuclei orientation (Fig. S2G). Other formins (Daam2, FHOD1, and FMN2) did not produce noticeable effect, or even slightly increased (INF2) these angles (Fig. S2G).

We further assessed the effect of the knockdowns of other major regulators of actin polymerization (*27*). Remarkably, the suppression of the actin-nucleating Arp2/3 complex via the knockdown of its major component ARPC2 (Fig. S4F), resulted in the inhibition of chirality (Fig. S4, A and B), whereas the overall effect of ARPC2 downregulation on the actin cytoskeleton appearance in these experiments was relatively mild and cell spreading was not impaired. The double knockdown of actin depolymerizing proteins, cofilins 1 and 2, also led to a pronounced inhibition of actin chirality (Fig. S4, C and D), while the knockdown of related protein, actin depolymerizing factor (ADF), did not diminish anti-clockwise actin cytoskeleton chirality (Fig. S4, D and E). Re-introduction of only cofilin-1 to the cofilins 1 and 2 depleted cells, was sufficient to restore anti-clockwise chirality of the cell (Fig. S4, D, E and G). Interestingly, unlike in the knockdown of mDia1, the knockdown of ARPC2 or cofilins only slightly reduced the degree of chiral orientation of microcultures on rectangles (Fig. S4H). Finally, the knockdowns of VASP and Mena, which are activators of actin filament elongation, did not affect the formation of asymmetric actin pattern in individual cells (Fig. S5, A to E) and chiral alignment of cells in microcultures (Fig. S5F).

### Knockdowns of actin monomers sequestering and capping protein, as well as treatment with actin monomers sequestering drugs reversed direction of actin cytoskeleton chirality

While depletion of mDia1, ARPC2, or cofilins reduced the asymmetry of actin pattern, the manipulations of several other actin associated proteins or actin pharmacological perturbations reversed the direction of chirality (Fig. 2). Profilins are abundant actin monomer sequestering proteins which can either augment or inhibit actin polymerization, depending on their biological context (*28*). We found that the siRNA-mediated knockdown of profilin 1 (Fig. S6A) reversed the direction of actin swirling in cells, resulting in sinistral chirality pattern (Fig. 2B and Movie S3). This effect can be rescued by the expression of exogenous mouse profilin 1 (Fig. 2, D and F). The curve showing the average tilt of radial fibers as a function of distance from the cell edge in profilin 1 siRNA cells was approximately a mirror-image of the curve for control cells (Fig. 2F). Cells lacking the less abundant profilin 2 (Fig. S6A) still demonstrated anti-clockwise chiral swirling of the actin cytoskeleton similar to control cells (Fig. 2, C and F). Capping proteins bind to the barbed end of actin filaments and interferes with both polymerization and depolymerization (*27*). Knockdown of CapZβ a subunit of capping protein CapZ, (Fig. S6B) resulted in the reversal of direction of radial fibers tilting so that sinistral actin pattern was formed (Fig. 2, E and F).

**Fig. 2.**
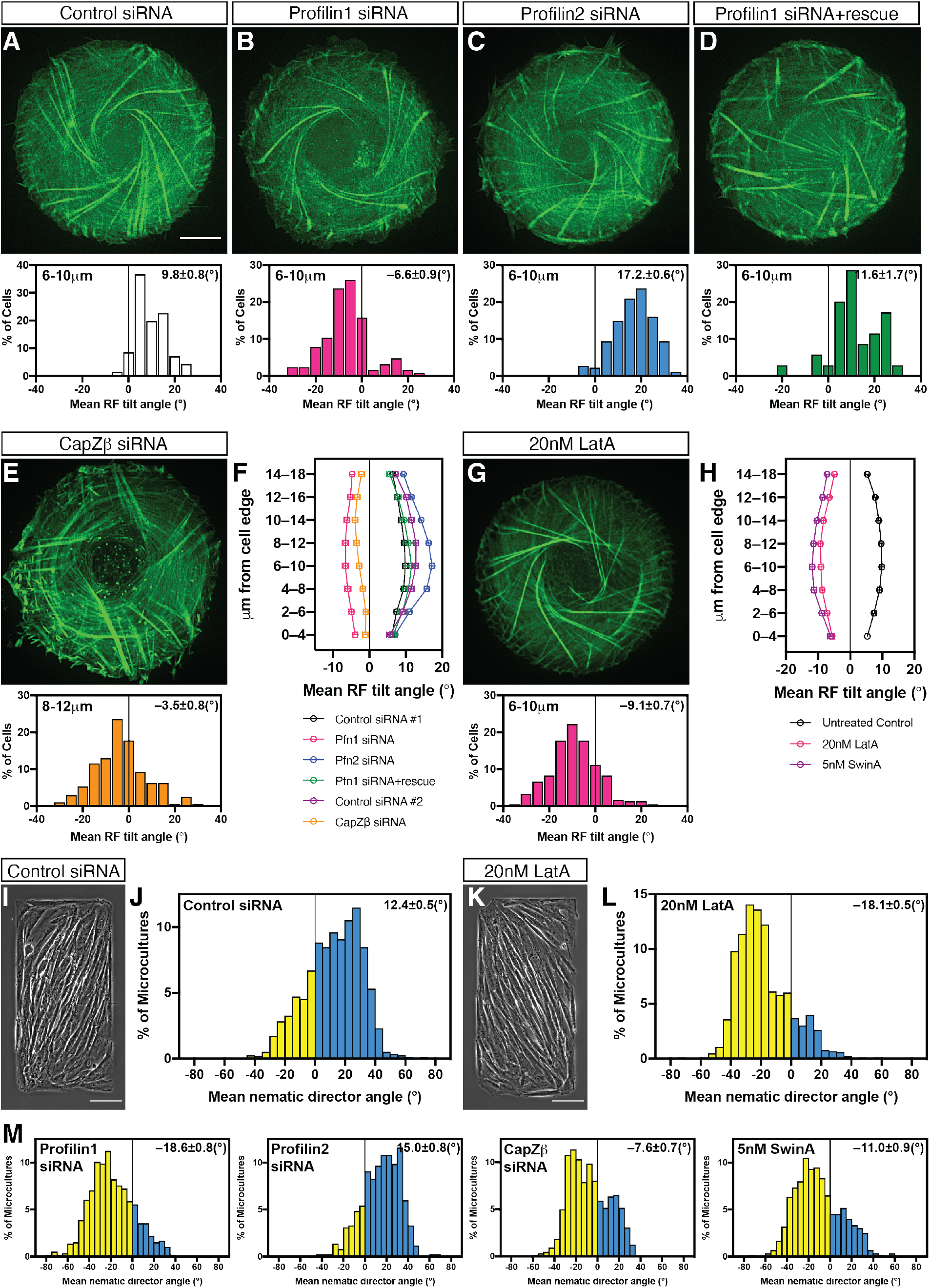
Genetic knockdowns and pharmacological treatments reversing the direction of the actin cytoskeleton chirality. (**A** to **D**) Typical examples of actin organization visualized by LifeAct-GFP in control siRNA (**A**), Profilin 1 (Pfn1) siRNA (**B**), Profilin 2 (Pfn2) siRNA (**C**), and Profilin 1 siRNA plus mouse Pfn1 full-length plasmid (**D**) transfected cells. The histograms (**A** to **D**) show the distribution of average RF tilt in the 6-10 μm annulus in cells under corresponding conditions. mean±*s.e.m* values of these average RF tilts were obtained from 71 control cells, 127 Profilin 1 knockdown cells, 182 Profilin 2 knockdown cells and 35 Profilin 1 knockdown cells rescued by co-transfection with full-length Profilin 1 plasmid. (**E**) Typical example of actin organization visualized by phalloidin-staining in CapZβ siRNA-transfected cells 6 hours after cell plating. Histogram under the image shows the distribution of average RF tilt in the 8-12 μm annulus. Mean±*s.e.m* value of average RF tilts was obtained from 208 cells. (**F**) Average values of RF tilts (mean±*s.e.m*) as a function of the distance from the cell edge for Profilins and CapZ² experiments. Control siRNA #1 and #2 represent the control cells used in experiments with Profilins and CapZβ respectively. (**G**) Typical example of actin organization visualized by phalloidin-labeling in cells treated with 20nM of latrunculinA (LatA). Histogram under image shows the distribution of average RF tilt in the 6-10 μm annulus. mean±*s.e.m* value of average RF tilts was obtained from 243 cells. (**H**) Average values of RF tilts (mean± *s.e.m*) as a function of the distance from the cell edge for untreated control cells (n= 274), 20nM LatA-treated cells (n= 243), and 5nM swinholideA (SwinA)-treated cells (n= 153). (**I** to **M**) Knockdowns of Profilin 1 and CapZβ and treatments with LatA and SwinA reversed the sign of cell orientation angle in microcultures confined to rectangular micropatterns. Phase-contrast image of control (**I**) and LatA-treated (**K**) cells 48 hours following plating on rectangular adhesive pattern. Histograms (**J**, **L** and **M**) show distributions of the values of mean nematic directors characterizing individual microcultures on rectangles for cells treated as indicated. The histograms were built based on average local cell orientation (nematic directors) values from 1168 control, 597 Profilin 1, 519 Profilin 2, and 661 CapZβ knockdown microcultures and 1031 LatA- and 602 SwinA-treated microcultures. See also Fig. S6. Mean±*s.e.m* values are indicated at the top right corner of each histogram. Scale bars, 10μm (**A** to **E** and **G**); 100μm (**I** and **K**). See also Movie S2.

Of note, even though the effects of knockdowns of CapZβ mDia1 and cofilins1&2 were reproducible in experiments where actin pattern was quantified in fixed phalloidin-stained cells, they were less pronounced in experiments where actin was visualized by expression of LifeAct fused with either GFP or mRuby fluorescent proteins. The reason for these discrepancies is unknown; it might be related to some side effects of LifeAct on actin polymerization (*29*).

By screening of actin polymerization affecting drugs, we found that latrunculin A, which sequester G-actin monomer and depolymerize F-actin (*30*), when applied at low concentration (20 nM), effectively reversed the direction of chirality – inducing sinistral actin pattern (Fig. 2G). The time course of chirality development in the presence of latrunculin A was the same as in control, while the average tilt of the radial fibers was equal in absolute value and opposite in direction as compared to control (Fig. 2H). The effect of latrunculin A was readily seen in both phalloidin-stained fixed cells and LifeAct-labeled live cells. Addition of latrunculin A reversed the direction of swirling in cells with an established anti-clockwise swirling (dextral) actin cytoskeleton (Movie S4). These effects were rapid and became evident within 1-2 hrs following latrunculin A addition. Washing out of latrunculin A resulted in rapid return to a “normal” anti-clockwise actin swirling (Movie S5). The fast effects of latrunculin A on chirality direction suggest that its action does not depend on any transcriptional effects. This is supported by additional evidence showing latrunculin A addition can reverse the direction of actin cytoskeleton chirality in enucleated cells (Fig. S7 and Movie S6). Among other actin polymerization affecting drugs, the actin filament severing and dimer forming drug swinholide A (*31*), at 5 nM, produced the same reversing effect on chirality direction as latrunculin A (Fig. 2H).

The genetic knockdowns and drug treatments that changed the direction of chirality of the actin cytoskeleton in individual cells induced corresponding changes in the chirality of alignment in microcultures on a rectangular pattern. Depletion of profilin 1 or CapZβ, or treatment with latrunculin A or swinholide A resulted in preferential cell alignment in the direction mirror-symmetrical to that of control cells (Fig. 2, I to M and S6C). In all these cases, the average orientation of multicellular groups in rectangles was tilted at negative angles relative to the long axis of the rectangle. Similar to results in individual cells, the knockdown of profilin 2 did not change the average direction of cell alignment as compared to control (Fig. 2M and S6C). Therefore, the effect of treatments affecting the direction of chirality in individual cells strongly correlates with their effect on the chirality of cell alignment in microcultures.

### Alteration of the chirality of individual cells and multicellular microcultures involves alpha-actinin1 crosslinking activity

Overexpression of actin crosslinking protein alpha-actinin1 resulted in the reduction of cell chirality and reversal of the direction of actin cytoskeleton swirling in a fraction of transfected cells in agreement with our previous study (*11*) (Fig 3, A and B and S8A). The effect of chirality reversal upon overexpression of alpha-actinin1 was also evident on multicellular microcultures confined to rectangular pattern (Fig. 3, C and D and S8E). Overexpression of alpha-actinin4 largely reduced the fraction of chiral cells, while overexpression of another actin crosslinker, Filamin A, had no effect (Fig. 3B and S8, B and C). Neither alpha-actinin1 knockdown alone nor the expression of truncated alpha-actinin construct (ABDdel-actinin) (*32*) which interfered with the actin crosslinking activity of alpha-actinin, changed the chirality direction in individual cells and cell groups (Fig. 3, E to I and M, and S8, D and E).

**Fig. 3.**
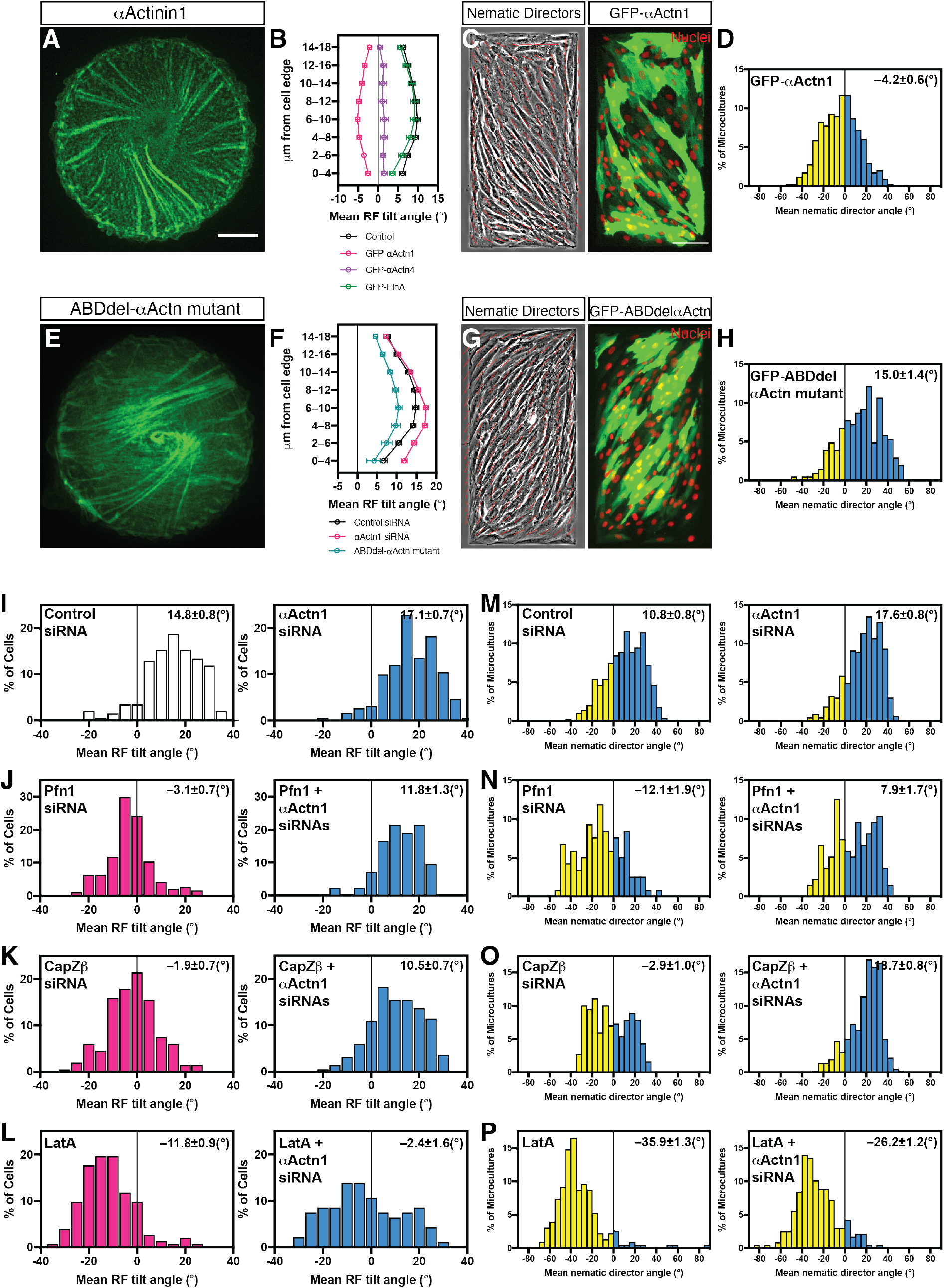
α-Actinin-1 is required for the reversal of chirality direction. (**A** to **D**) Effects of overexpression of α-actinin-1 and other crosslinking proteins on radial fiber (RF) tilt and cell alignment in microcultures. (**A**) Image of GFP-α-actinin-1 expressing cell. α-actinin-1-positive RFs tilt follows clockwise swirling. (**B**) Average values of RF tilts (mean±*s.e.m*) as a function of the distance from the cell edge for control cells (n= 70), and cells overexpressing α-actinin-1 (GFP-αActn1; n= 156), α-actinin-4 (GFP-αActn4; n= 92) and filamin A (GFP-FlnA; n= 69). (**C** and **D**) Effect of GFP-α-actinin-1 overexpression on the sign of cell orientation angle in microcultures confined to rectangular micropatterns. (**C**) Phase-contrast image overlayed with local nematic directors (red lines) (left) and distribution of GFP-αactinin-1 transfected cells in the same field (right). The nuclei were labelled with Hoechst 33342 (pseudo-colored red). (**D**) Histogram showing distribution of the angles of mean nematic directors characterizing individual microcultures for GFP-α-actinin-1 overexpressing cells. The histogram was built based on 745 microcultures. (**E** to **H**) Effects of inhibition of α-actinin-1 crosslinking function on RF tilt and cell alignment in microcultures. (**E**) Actin organization in GFP-ABDdel-α-actinin mutant expressing cell visualized by mRuby-LifeAct (pseudo-colored green). (**F**) Average values of RF tilts (mean±*s.e.m*) as a function of the distance from the cell edge for control siRNA-transfected cells (n= 203), α-actinin-1 siRNA-transfected cells (n= 264) and GFP-ABDdel α actinin mutant expressing cells (n= 85). (**G** and **H**) Effect of GFP-ABDdel-α-actinin mutant overexpression on the sign of cell orientation angle in microcultures. (**G**) Phase-contrast image overlayed with local nematic directors (red lines) (left) and distribution of GFP-ABDdel-α-actinin mutant transfected cells in the same field (right). The nuclei were labelled with Hoechst 33342 (pseudo-colored red). (**H**) Histogram showing distribution of the angles of mean nematic directors characterizing individual microcultures for GFP-ABDdel-αactinin mutant overexpressing cells. The histogram was built based on 206 microcultures. (**I** to **L**) The reversion of RF tilt is α-actinin1-dependent. The histograms show the distribution of average RF tilt in the 6-10 μm (**I**, **J**, **L**) or 8-12 μm (**K**) annulus for (**I**) control siRNA-transfected cells (n= 203) and α-actinin-1 siRNA-transfected cells (n= 192), (**J**) Profilin 1(Pfn1) siRNA-transfected cells (n= 194) and Pfn1α-actinin-1 siRNAs-transfected cells (n= 42), (**K**) CapZβ siRNA-transfected cells (n= 201) and CapZβ-actinin-1 siRNAs-transfected cells (n= 219), and (**L**) 20nM LatA-treated cells (n= 75) and α-actinin-1 siRNA-transfected cells treated with 20nM LatA (n= 94). (**M** to **P**) The reversion of the sign of cell alignment angle in microcultures by Profilin 1 and CapZβ knockdowns, but not by LatA treatment, depends on α-actinin-1. Histograms showing distributions of the angles of mean nematic directors characterizing the microcultures for (**M**) control siRNA-transfected cells (n= 499) and α-actinin1 siRNA-transfected cells (n= 430), (**N**) Pfn1 siRNA-transfected cells (n= 118) and Pfn1α-actinin-1 siRNAs-transfected cells (n= 135), (**O**) CapZβ siRNA-transfected cells (n= 369) and CapZβ-actinin-1 siRNAs-transfected cells (n= 360), and (**P**) 20nM LatA- treated cells (n= 230) and α-actinin-1 siRNA-transfected cells treated with 20nM LatA (n= 237). Mean±*s.e.m* values are indicated at the top right corner of each histogram. Scale bars, 10μm (**A** and **E**); 100μm (**C** and **G**).

In view of the involvement of alpha-actinin1 in reversal of cell chirality in these and other study (*23*), we further investigated the combined effect of alpha-actinin1 loss of function and experimental manipulations which reversed chirality direction. The double knockdowns of profilin 1 and alpha-actinin1 (Fig. 3J), CapZβ and alpha-actinin1 (Fig. 3K), and latrunculin A treatment of alpha-actinin1 knockdown cells or ABDdel-actinin expressing cells (Fig. 3L and S8F), all resulted in dextral anti-clockwise chirality comparable with that in control cells (Fig. 3I). Thus, reversal of chirality by depletion of either profilin 1 or CapZβ, or latrunculin treatment requires alpha-actinin1 function. In microcultures, knockdown of alpha-actinin1 similarly prevents the reversal of chirality upon profilin 1 (Fig. 3N) or CapZβ(Fig. 3O) depletion, even though not upon latrunculin A treatment (Fig. 3P).

Since several studies have clearly demonstrated the importance of myosin 1c and 1d in the chiral morphogenesis of Drosophila organs (*5, 6, 18*), we checked whether these myosins are involved in individual and collective chirality development in our experimental systems. We were not able to detect any effects of myosin 1c or myosin 1d knockdown on the development of the chiral actin pattern in individual cells or in asymmetric alignment of cells in microcultures (Fig. S9). Thus, either small residual amount of these myosins is sufficient for chirality development or these myosins are not involved in chirality development in human fibroblasts.

In order to assess whether any of the knockdown effects observed above could be attributed to an associated alteration in the expression level of other proteins involved in chirality regulation, we examined the transcriptional profile of cells depleted of major proteins strongly associated with chirality phenotype (Fig. S10). We found that, with few exceptions (Fig. S10), knockdowns of the members of this group of proteins (mDia1, ARPC2, cofilins 1&2, CapZβ, profilin 1, and alpha-actinin1) only slightly, if at all, affected the expressions of other members of the group. These data suggested that phenotypic changes observed upon knockdown of these proteins are not mediated by transcriptional regulation of the expression of other members of this group. This is consistent with the apparent transcription-independent effect on chirality observed in latrunculin A-treated enucleated cells, as mentioned above (Fig. S7).

### Correlation between chirality of the actin cytoskeleton in individual cells and chiral alignment of cells in multicellular microcultures

Altogether, in our experiments, we characterized the effects of 35 different pharmacological and genetic manipulations on the emergence of left-right actin cytoskeleton asymmetry in individual confined cells, as well as on chiral cell alignment of confined multicellular groups. To analyze the interrelationship between the establishment of left-right asymmetry in these two systems, we plotted the average angle between the nematic directors characterizing the alignment of cells in cell groups versus the average tilt of radial fiber segments located between 6 – 10 microns from the cell edge (Fig. 4). In spite of some discrepancies, as mentioned earlier, the correlation between these two parameters was highly significant (Fig. 4 and S11 and Table S1). Altogether these data clearly demonstrate the role of actin cytoskeleton asymmetry in individual cells in the establishment of collective asymmetry of cell alignment in cell groups.

**Fig. 4.**
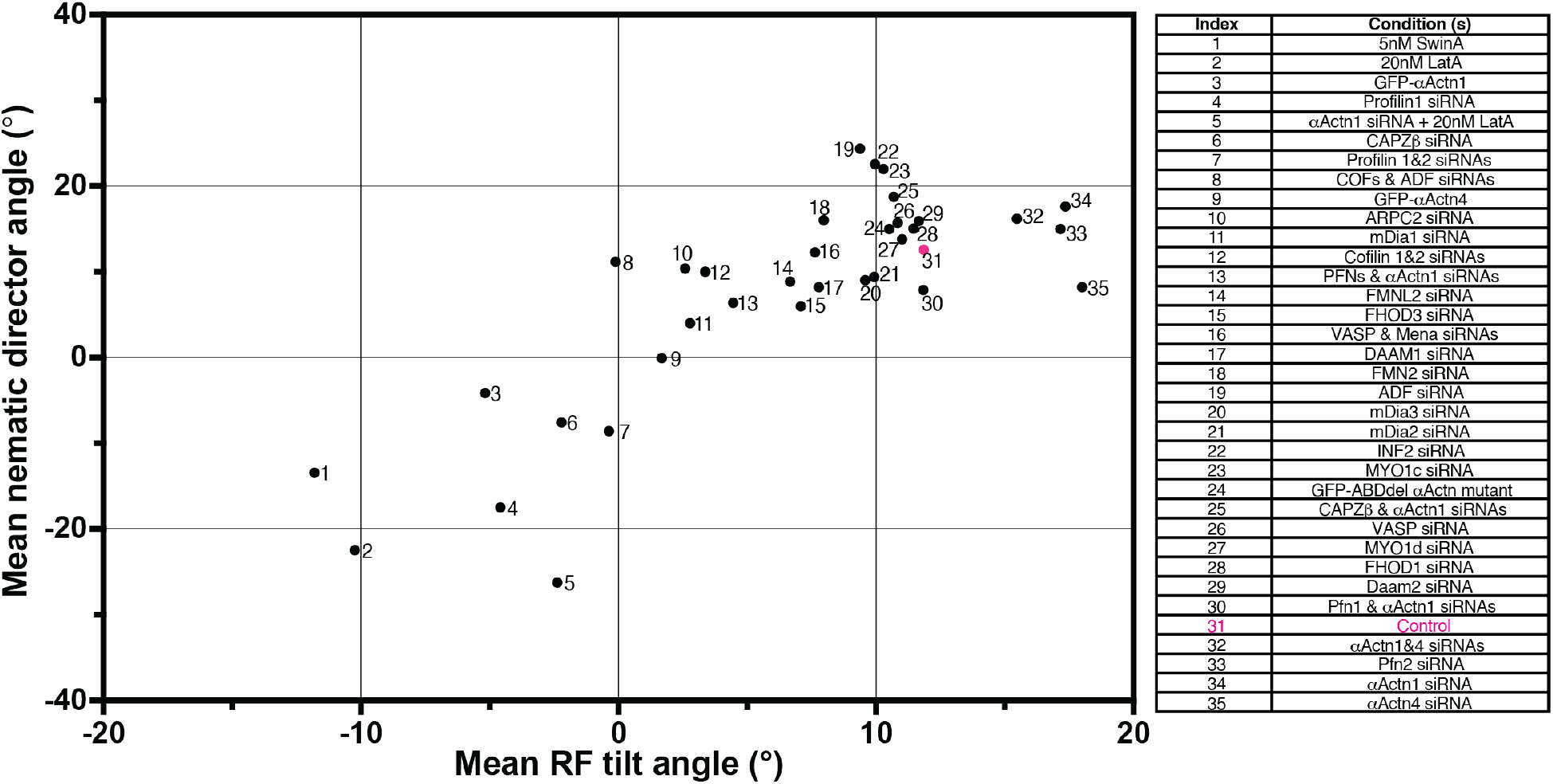
The correlation between mean radial fiber tilt angle in individual cells and mean nematic director angle in microcultures. Mean nematic director angles for rectangular microcultures (*y*-axis) are plotted against mean radial fiber tilt angles at the 6-10 μm annulus (*x*-axis) for each type of treatment. Each dot represents average data from pooled experiments under respective conditions as indicated in the list on the right. The data are ordered and indexed according to ascending radial fiber tilt angles (*x*-axis). Position of control cell is in red. Pearson correlation coefficient, r = 0.8104, *\*\*\*\*p* <0.0001. Numbers of cells and microcultures analyzed and the values of the means±*s.e.m* can be found in Table S1. See also Fig. S11.

## Discussion

The key improvement which made this study possible was the development of rigorous quantitative methods which permitted us to perform a large-scale assessment of the degree of left-right asymmetry in the organization of the actin cytoskeleton in individual cells and the alignment of cells in confined cell groups. The process of left-right asymmetric actin swirling in isotropic discoid cells is manifested by unilateral tilting of the radial fibers. Thus, using deep-learning computational image analysis, we determined the angles characterizing the degree of deviation of these fibers from the radial direction in individual cells. Formation of confluent cell monolayer in microcultures confined to a rectangular micropattern resulted in development of a prevalent angle of cell alignment. We assessed the deviation between cell alignment axis and the long axis of the rectangle by measuring either the average angle of local nematic directors in phase contrast images or the average angle of long axes of elliptical cell nuclei. These objective measurements of the left-right asymmetry in individual cells and cell groups allowed us to make quantitative comparisons between the processes of asymmetric actin cytoskeleton organization and asymmetric cell alignment.

In contrast to earlier studies that focused on a single gene as the main regulator of left-right asymmetry, our study revealed that multiple actin-associated proteins are involved in the control of chiral morphogenesis in individual cells and multicellular microcultures. Among the proteins involved in actin assembly, not only formins but also Arp2/3 complex, cofilins, capping protein and profilin appeared to be potent regulators of left-right asymmetry development. Of note, all of these proteins were shown to function in cooperation or competition with formins, in particular mDia1 (*33–37*). In addition, major crosslinking protein alpha-actinin1 was shown to function as a co-factor for chirality changes induced by the regulators of actin polymerization. We have not however, found a noticeable effect of downregulation of myosin 1c or 1d on left-right asymmetry development in individual cells or cell groups, yet these motor proteins are known to be strong regulators of the left-right asymmetry in the development of *Drosophila* organs (*5, 6, 18*). Altogether, these and other data (*14*) suggest that the translation of actin filament asymmetry into cellular and tissue asymmetric morphogenic behavior could be species-, and even cell-type-specific. The asymmetry emerging from actin chiral helical organization could be a common denominator of left-right asymmetry development in many systems and it cannot be excluded that some other actin regulating proteins, in addition to those identified in this study, will be found to be involved.

A remarkable type of response observed in our study was switching from dextral to sinistral chirality. Reversal of the direction of chirality is a phenomenon observed at the organismal level, which include *situs inversus* in vertebrates, reversed chirality of hindgut in flies, and sinistral chirality in snails (*1–3*). In our studies, we found that the actin cytoskeleton of individual cells can also demonstrate a pattern of organization that looks like a mirrored reflection of the normal chiral pattern. The most striking examples are knockdowns of profilin 1 and CapZβ subunit of capping protein CapZ, which both led to negative average tilting of the radial fibers in individual cells and in the orientation of alignment of cell groups. Another group of treatments that efficiently reversed both individual and collective chirality direction was treatment with low concentrations of actin polymerization inhibitors, latrunculin A and swinholide A. We showed previously that latrunculin A promoted clockwise actin swirling in initially radially symmetrical epithelial cells (*14*). The factors promoting sinistral chirality appeared to be specific – depletion of profilin 1 but not profilin 2, or treatment with latrunculin A/swinholide A but not other polymerization inhibitors (*14*). While a unifying mechanism underlying the reversal of swirling direction remains to be elucidated, these data suggest that cellular handedness can be reversed by some specific perturbation of actin polymerization.

A new type of regulatory signature revealed in this study is the requirement for alpha-actinin function. Knockdown of alpha-actinin1 changed the organization of actin cytoskeleton in individual cells (*11, 38*) but did not interfere with normal dextral chirality in individual cells ((*11*) and present study) and cell groups. However, the presence of alpha-actinin1 appeared to be critical for sinistral (clockwise) asymmetry of the actin cytoskeleton. Combination of knockdowns of either profilin 1 or CapZβ with knockdown of alpha-actinin1, abolished the reversal of chirality direction in individual cells and cell groups, suggesting that alpha-actinin1 is needed for reversal of cell chirality. Alpha-actinin1 is a major crosslinking protein in radial fibers and its function in these processes may depend on its possible role in hindering individual filament rotation (in line with a model in (*11*)) or its possible involvement in radial fiber twisting.

Our experimental systems allowed us to perform systematic quantitative comparison between effects of diverse genetic and pharmacological treatments on development of left-right asymmetry in individual cells and multicellular microcultures confined to rectangular patterns. Asymmetric alignment of cells in our microcultures resembles chiral behavior seen in cells confined to stripes or ring-shaped patterns (*8–10*). Our study revealed a remarkable correlation between responses of individual cells and cell collectives in microcultures. With only few exceptions, the treatments which affected formation of asymmetric actin pattern in individual cells also affected asymmetric alignment of cell groups. Treatments that reversed actin chirality direction in individual cells always resulted in a change of the direction of average cell alignment in cell groups. These data, in line with (*39*), provide strong experimental support to the hypothesis that development of chiral organization in multicellular cultures, tissues, and organs is determined by chirality of the actin cytoskeleton in the individual cell.

The hypothesis of the role of asymmetric self-organization of the actin cytoskeleton in emergence of chirality in tissues, organs, and even in whole organism could unify the existing data on the development of left-right asymmetry in snails (*19, 20*) and in *Drosophila* (*5, 6, 18, 21*), and probably for asymmetric heart looping in vertebrate development (*4*). The formation of asymmetry of heart and visceral organs positioning in vertebrate deserves special discussion. It is well-established that in many species the key asymmetric factor triggering the signaling cascade determining left-right body asymmetry is an asymmetric flow generated by ciliated cells in the node (left-right organizer of the embryo) (*1, 2*). Recent studies have shown, however, that in birds and reptilia the nodal asymmetry does not depend on cilia (*1*). Thus, the cilia may function as an amplifier, but not the primary source of asymmetry. Since the position and orientation of basal bodies can in principle be regulated by the actin cytoskeleton (*40*), this suggests that the primary asymmetric factor in nodal cilia-dependent systems is could still be the intrinsic asymmetry of the actin cytoskeleton. Extensive future studies are necessary to explore this possibility. In conclusion, our study revealed an actin polymerization-dependent mechanism of establishment of left-right asymmetry in individual cells and cell groups which could be involved in the development of left-right asymmetry in organs and organisms.

## Acknowledgments

We thank M. M. Kozlov (Tel Aviv University, Israel) and T. Hiraiwa (MBI, Singapore) for discussion, T. B. Saw (MBI, Singapore) for reviewing the manuscript, M. Davidson (The Florida State University, Tallahassee, USA), P. Roca-Cusachs, M. Pan, M. Sheetz and C. G. Koh for providing reagents, A. Wong (MBI, Singapore) for expert help in paper editing and the microscopy and nanofabrication core facilities at the Mechanobiology Institute for technical help.

## Funding

The research is supported in part by the Singapore Ministry of Education Academic Research Fund Tier 2 (MOE Grant No: MOE2018-T2-2-138, MOE2019-T2-1-099, MOE2019-T2-02-014), and Tier 3 (MOE Grant No: MOE2016-T3-1-002), the National Research Foundation, Prime Minister’s Office, Singapore, and the Ministry of Education under the Research Centers of Excellence program through the Mechanobiology Institute, Singapore (ref no. R-714-006-006-271), and by the Singapore Ministry of Health’s National Medical Research Council under its “Open Fund - Young Individual Research Grant” (Grant No: OFYIRG18may-0041).

## Author contributions

YHT and ADB designed the experiments. YHT, WJG and XY performed most experiments. JH, IYYT, SS, SJ, VT, SFHB, and WH contributed to some experiments. WJG, XY and HTO developed image analysis tools. YHT and ADB wrote the manuscript with input from all of the authors.

## Competing interests

Authors declare no competing interests.

## Data and materials availability

All data are available in the main text or the supplementary materials. All image analysis codes are available upon request.

## List of Supplementary Materials

Materials and Methods

Figs. S1 to S11

Tables S1 to S3

References (41–47)

Movies S1 to S6

## Supplementary Figures

**Fig. S1.**
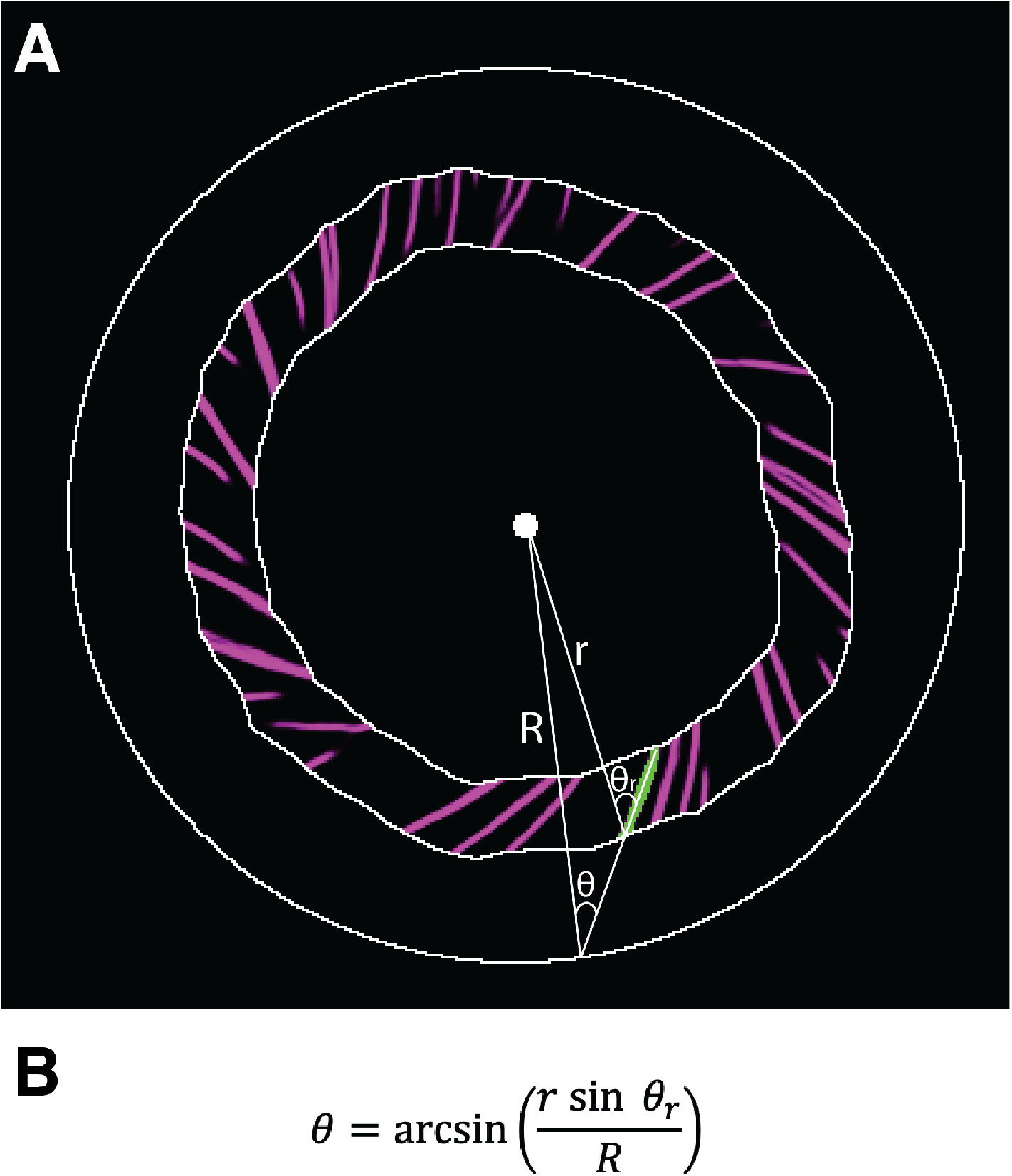
Calculation of radial fiber tilt. The tilts of all radial fiber segments in concentric belts (annuli) located at given distance from the cell edge were measured. The radial fiber segments in the 6-10 μm annulus is shown in (**A**). Measurement of the tilt of a single radial fiber segment (highlighted in green) is illustrated in (**A**) and calculated according to the formula in (**B**). 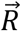 connects the cell centroid and intersection of the continuation of the radial fiber segment with the edge of the cell. 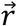 connects the cell centroid and intersection of radial fiber with outer edge of the annulus. θ and θ_r_ are the angles between the radial fiber segment and 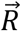 and 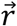 respectively.

**Fig. S2.**
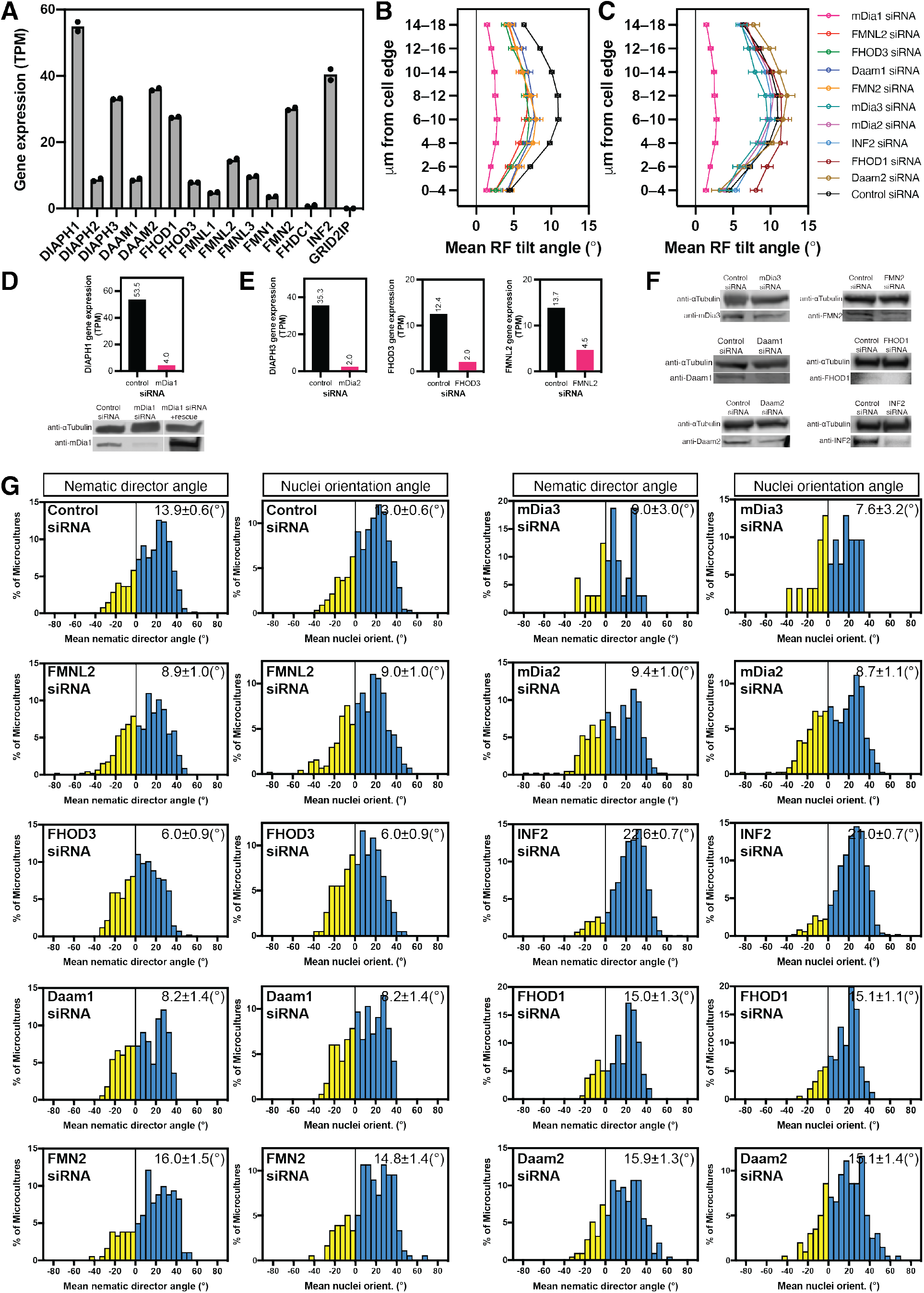
Effect of knockdown of formin family proteins on left-right asymmetry of actin organization in individual cells and chiral cell alignment in microcultures. (**A**) Transcriptome profiling of gene expression levels (transcripts per million; TPM) of the 15 mammalian formin members identified by RNA-sequencing (RNA-seq) (mean values of n= 2 experiments) in human fibroblasts. (**B** and **C**) Average values of radial fiber (RF) tilts (mean±*s.e.m*) as a function of the distance from the cell edge for formin family members, knockdown of which slightly reduced (**B**) or did not apparently affect (**C**) actin cytoskeleton chirality. Graphs corresponding to mDia1 (red) and control (black) siRNAs are presented in both (**B**) and (**C**). Mean±*s.e.m* of the distribution of average RF tilt in the 6-10 μm annulus of the various knockdowns can be found in Table S1. (**D**) siRNA knockdown of mDia1 (DIAPH1) in fibroblasts as verified by RNA-profiling (top) and western blot (bottom). Rescue of mDia1 knockdown cells by co-transfection with mDia1 full-length plasmid is shown in lane 3 of western blot. (**E** and **F**) Gene expression levels (**E**) and western blots (**F**) showing individual formin protein levels in control siRNA and formin specific siRNAs-treated cells. α-Tubulin was used as loading controls in (**D**) and (**F**). (**G**) Quantification of chiral alignment of cells with formin protein knockdowns in microcultures as characterized by mean nematic directors or mean nuclei orientation. Mean±*s.e.m* values are indicated at the top right corner of each histogram. Sample sizes (n) for (**B**), (**C**) and (**G**) can be found in Table S1.

**Fig. S3.**
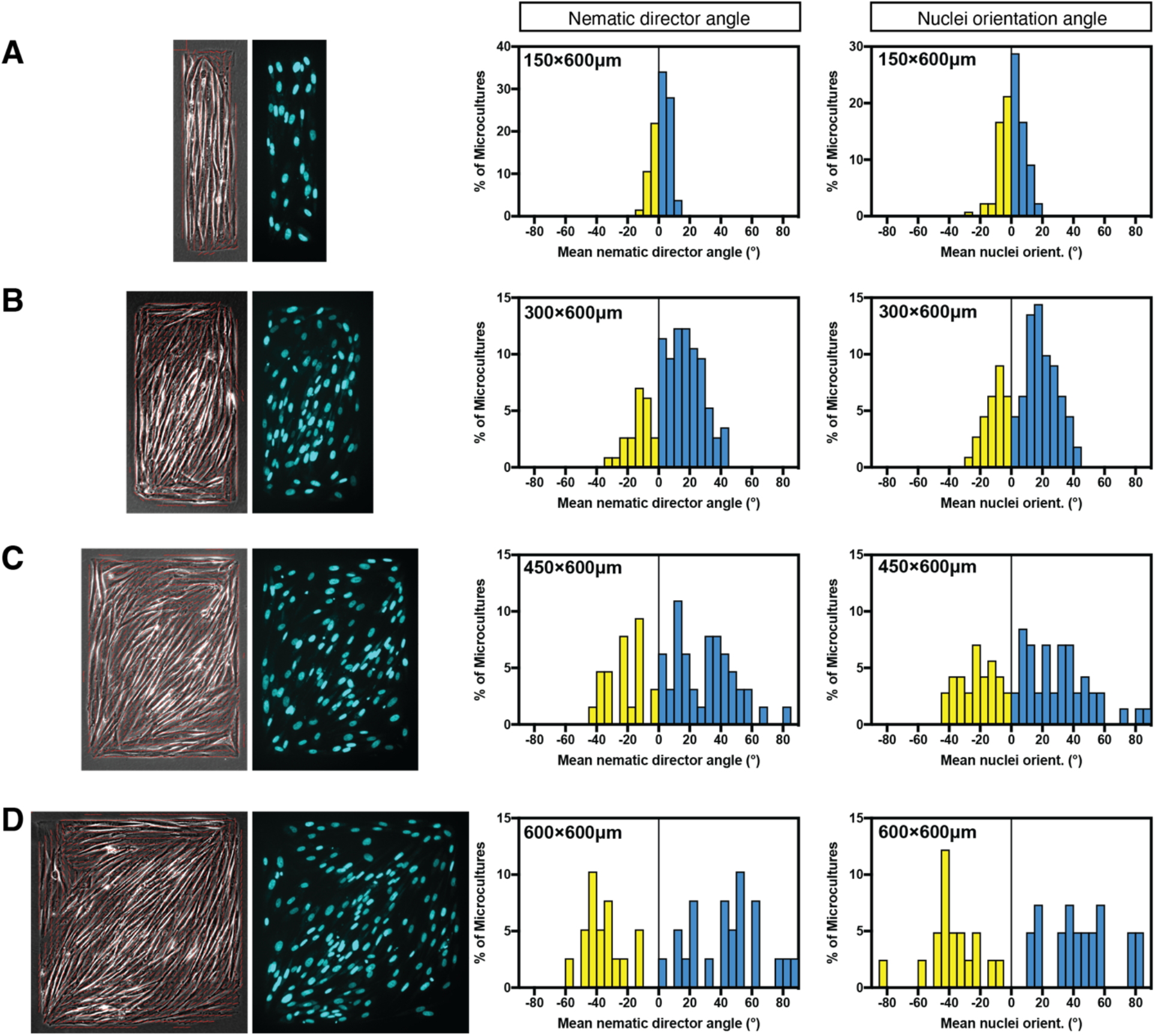
Relationship between aspect ratio of rectangular micropattern and left-right asymmetric cell alignment. Phase-contrast image overlayed with local nematic directors (red lines) (left) and the corresponding image of cell nuclei stained with Hoechst 33342 (right) of microcultures on 150×600 (**A**), 300×600 (**B**), 450×600 (**C**) and 600×600 (**D**) μm rectangular micropatterns. (**A** to **D**) Histograms showing distributions of the values of mean nematic directors and mean nuclei orientation characterizing individual microcultures on rectangles in respective conditions. The histograms were built based on average local cell orientation (nematic directors) values from 132 (**A**), 114 (**B**), 64 (**C**) and 39 (**D**) microcultures, or average nuclei orientation values from 132 (**A**), 111 (**B**), 71 (**C**) and 41 (**D**) microcultures respectively.

**Fig. S4.**
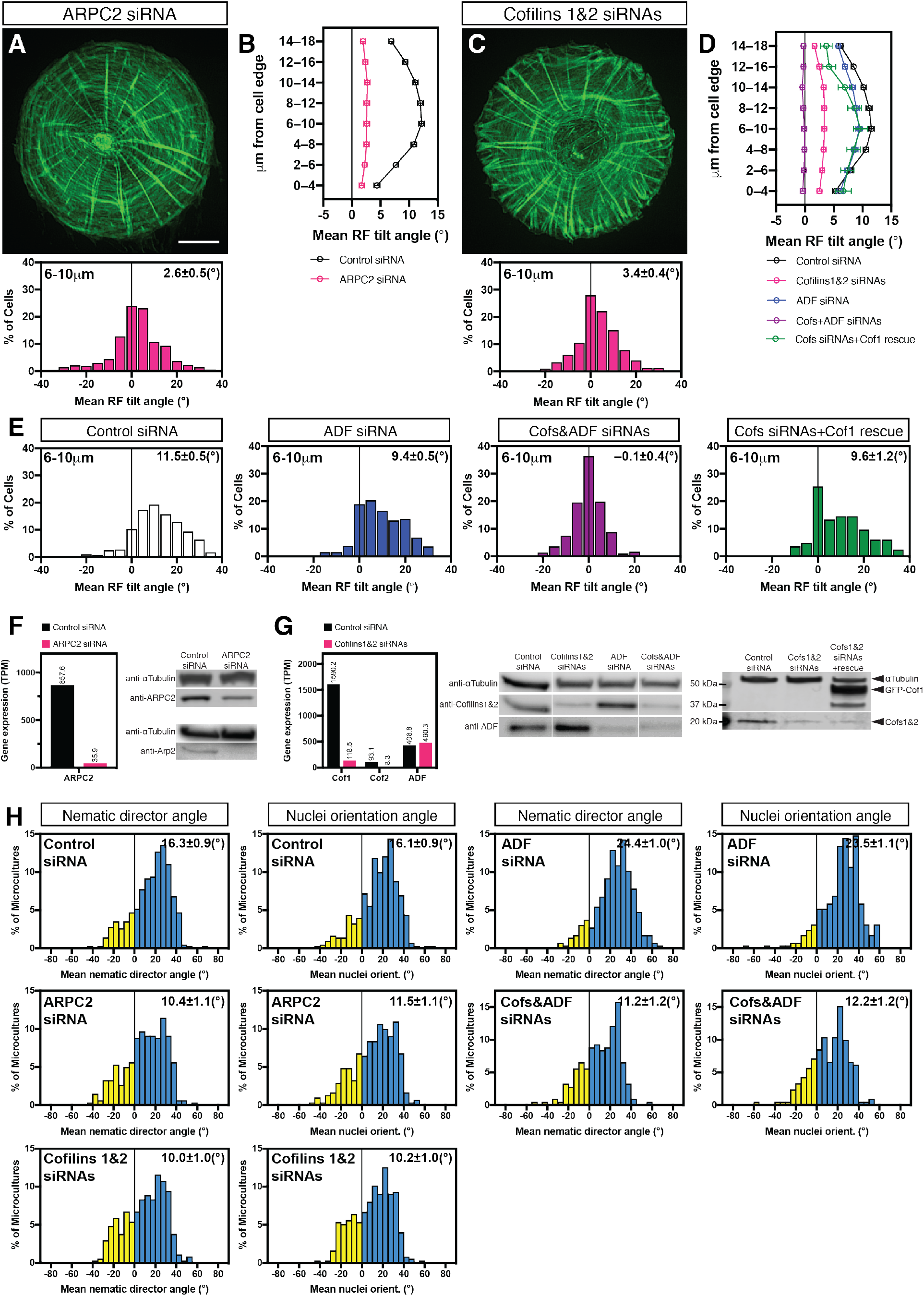
Quantification of left-right asymmetry in actin organization and chiral alignment of cells for knockdown of ARPC2 and actin-depolymerization factor (ADF)/cofilin family proteins. Actin organization visualized by phalloidin staining in ARPC2 siRNA (**A**) and Cofilins 1 and 2 siRNAs (**C**) transfected cells 6 hours following cell plating on circular pattern. The histograms (**A** and **C**) show the distribution of average radial fiber (RF) tilt in the 6-10 μm annulus in cells under corresponding conditions. (**B** and **D**) Average values of RF tilts (mean±*s.e.m*) as a function of the distance from the cell edge for experiments with ARPC2 (**B**) and ADF/Cofilins (**D**) respectively. (**E**) The histograms showing the distribution of average RF tilt in the 6-10 μm annulus in cells transfected as indicated. (**F**) siRNA knockdown of ARPC2 in fibroblast as verified by RNA-profiling (left) and western blot (right upper). Level of Arp2 is also reduced in ARPC2 siRNA transfected cells as compared to control cells (right lower). (**G**) siRNA knockdown of ADF/Cofilins family proteins as verified by RNA-profiling (left) and western blot (center). Rescue of Cofilins 1&2 knockdown cells by co-transfection with GFP-Cofilin 1 full-length plasmid is shown in lane 3 of western blot (right). (**H**) Quantification of chiral alignment of cells in microcultures under corresponding conditions as characterized by mean nematic directors or mean nuclei orientation. Mean±*s.e.m* values are indicated at the top right corner of each histogram. Scale bars, 10μm (**A** and **C**). Sample sizes (n) for (**A**), (**C**), (**E**) and (**H**) can be found in Table S1.

**Fig. S5.**
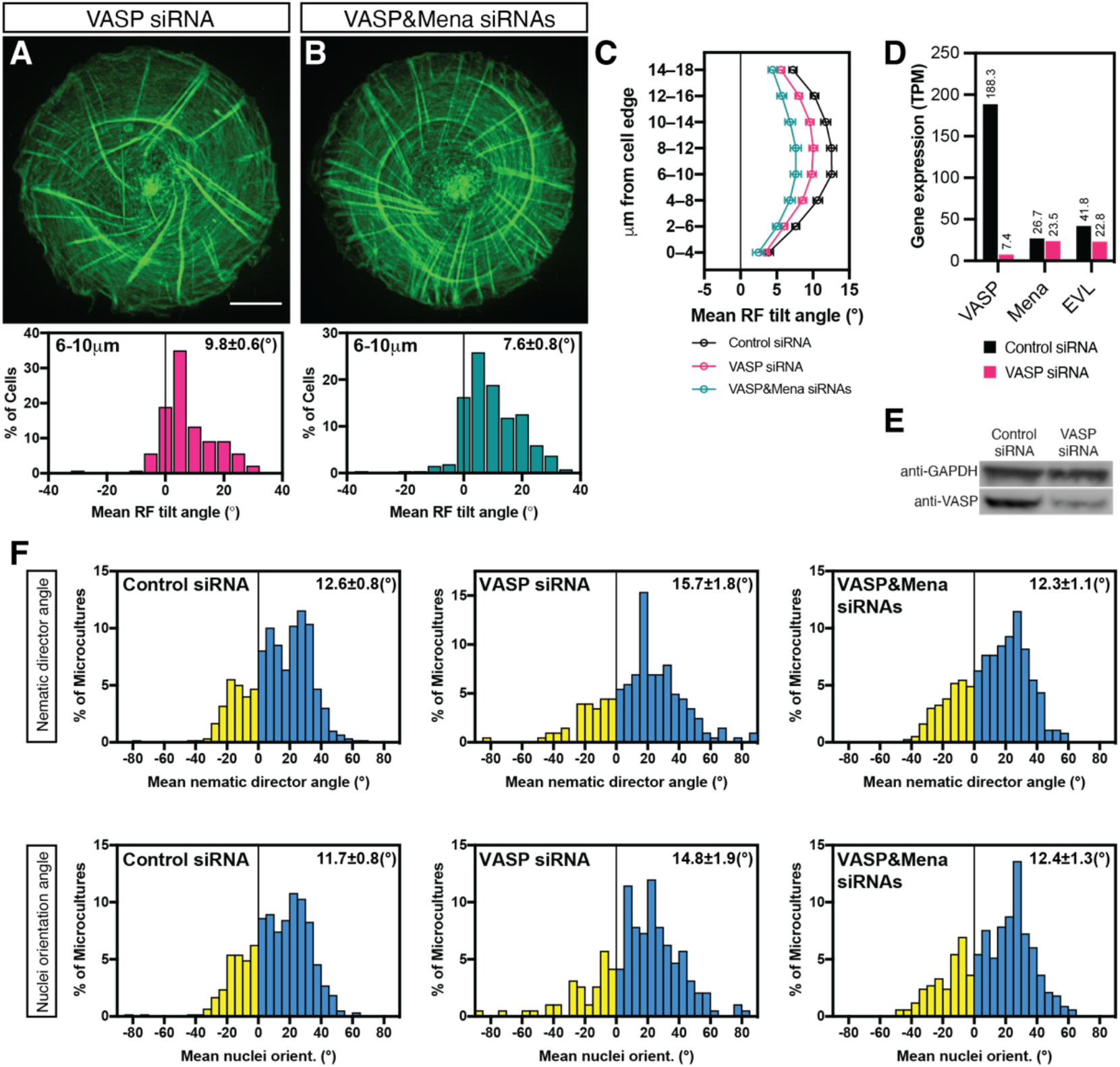
Effect of knockdown of actin filament elongator VASP on left-right asymmetry of actin organization in individual cells and chiral cell alignment in microcultures. Actin organization visualized by phalloidin staining in VASP siRNA (**A**) and VASP and Mena siRNAs (**B**)transfected cells 6 hours following cell plating on circular pattern are shown. The histograms show the distribution of average RF tilt in the 6-10 μm annulus in cells under corresponding conditions. (**C**) Average values of RF tilts (mean±*s.e.m*) as a function of the distance from the cell edge. Mean±*s.e.m* values of these average RF tilts (in **A** to **C**) were obtained from 214 control cells, 271 VASP knockdown cells and 143 VASP and Mena knockdown cells. (**D**) Transcriptome profiling of gene expression levels (transcripts per million; TPM) of the Ena/VASP family proteins by RNA-sequencing in control- and VASP- siRNA transfected human fibroblasts (n=1 experiment). (**E**) Western blot showing VASP level in cells treated with scramble (control) or anti-VASP siRNA; GAPDH was used as a loading control. (**F**) Histograms showing distributions of the values of mean nematic directors (upper row) and mean nuclei orientation (lower row) characterizing individual microcultures on rectangles. The histograms and mean±*s.e.m* values were built based on average local cell orientation (nematic directors) values from 598 control, 202 VASP knockdown and 366 VASP and Mena knockdown microcultures, or average nuclei orientation values from 593 control, 192 VASP and 331 VASP and Mena knockdown microcultures respectively. Scale bars, 10μm (**A** and **B**).

**Fig. S6.**
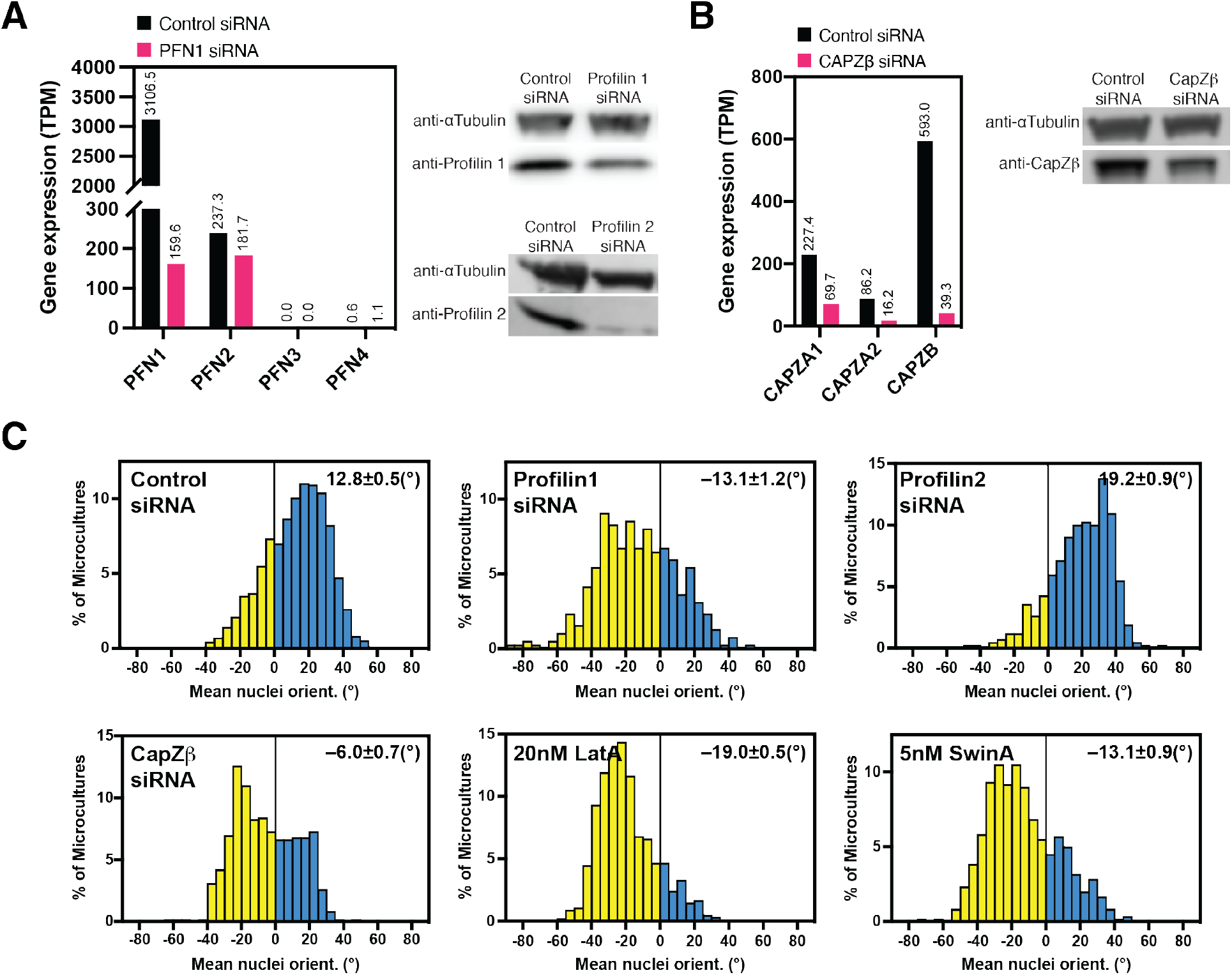
Knockdown of Profilin 1 and CapZβ protein and their effect on nuclei orientation of cells in microcultures. (**A**) Transcriptome profiling of gene expression levels (transcripts per million; TPM) of Profilin 1-4 isoforms identified by RNA-sequencing (mean values of n= 2 experiments) in control siRNA and Profilin 1 siRNA transfected cells. Western blot showing Profilin 1 (upper blot) and Profilin 2 (lower blot) levels in cells treated with scramble (control), Profilin 1 siRNA or Profilin 2 siRNA; α-tubulin was used as a loading control. Rescue of Profilin 1 knockdown cells by co-transfection with Pfn1-P2A-eGFP full-length plasmid is shown in lane 3 of western blot (upper blot). (**B**) Transcriptome profiling of gene expression levels of CapZA1, CapZA2 and CapZB identified by RNA-sequencing in control siRNA and CAPZβ siRNA transfected cells (n= 1 experiment). Western blot showing CAPZβlevels in cells treated with scramble (control) or CAPZβsiRNA; α-tubulin was used as a loading control. (**C**) Quantification of chiral alignment of cells in microcultures under respective conditions as characterized by mean nuclei orientation (correspond to Fig. 2, **J**, **L** and **M**). The histograms were built based on average nuclei orientation values from 1144 control, 386 Profilin 1 knockdown, 420 Profilin 2 knockdown, 620 CapZβ knockdown microcultures and 1031 LatA- and 601 SwinA-treated microcultures. Mean±*s.e.m* values are indicated at the top right corner of each histogram.

**Fig. S7.**
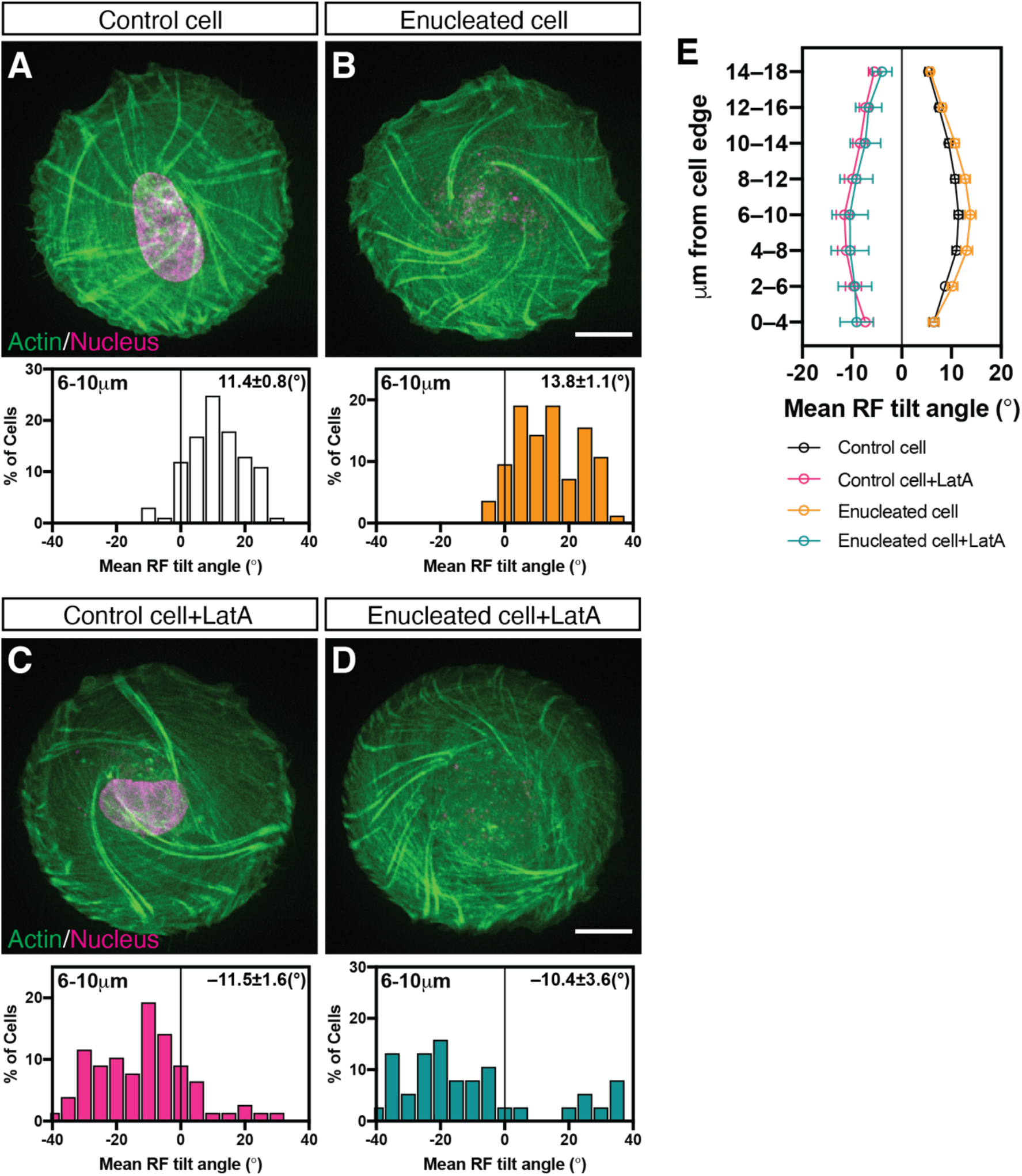
Evaluation of chirality in enucleated and latrunculin A-treated enucleated cells. Actin organization (green) as visualized by LifeAct-GFP and nuclei (pseudo-colored magenta) as labelled by Hoechst 33342, in control cell (**A**), enucleated cell (**B**), control cell treated with 20nM LatA (**C**) and enucleated cell treated with 20nM LatA (**D**). The histograms (**A** to **D**) show the distribution of average RF tilt in the 6-10 μm annulus in cells under corresponding conditions. (**E**) Average values of RF tilts (mean±*s.e.m*) as a function of the distance from the cell edge. Mean±*s.e.m* values of these average RF tilts (in **A** to **E**) were calculated from 101 control cells, 84 enucleated cells, 78 LatA-treated control cells and 38 LatA-treated enucleated cells. Scale bars, 10μm (**A** to **D**). See also Movie S6.

**Fig. S8.**
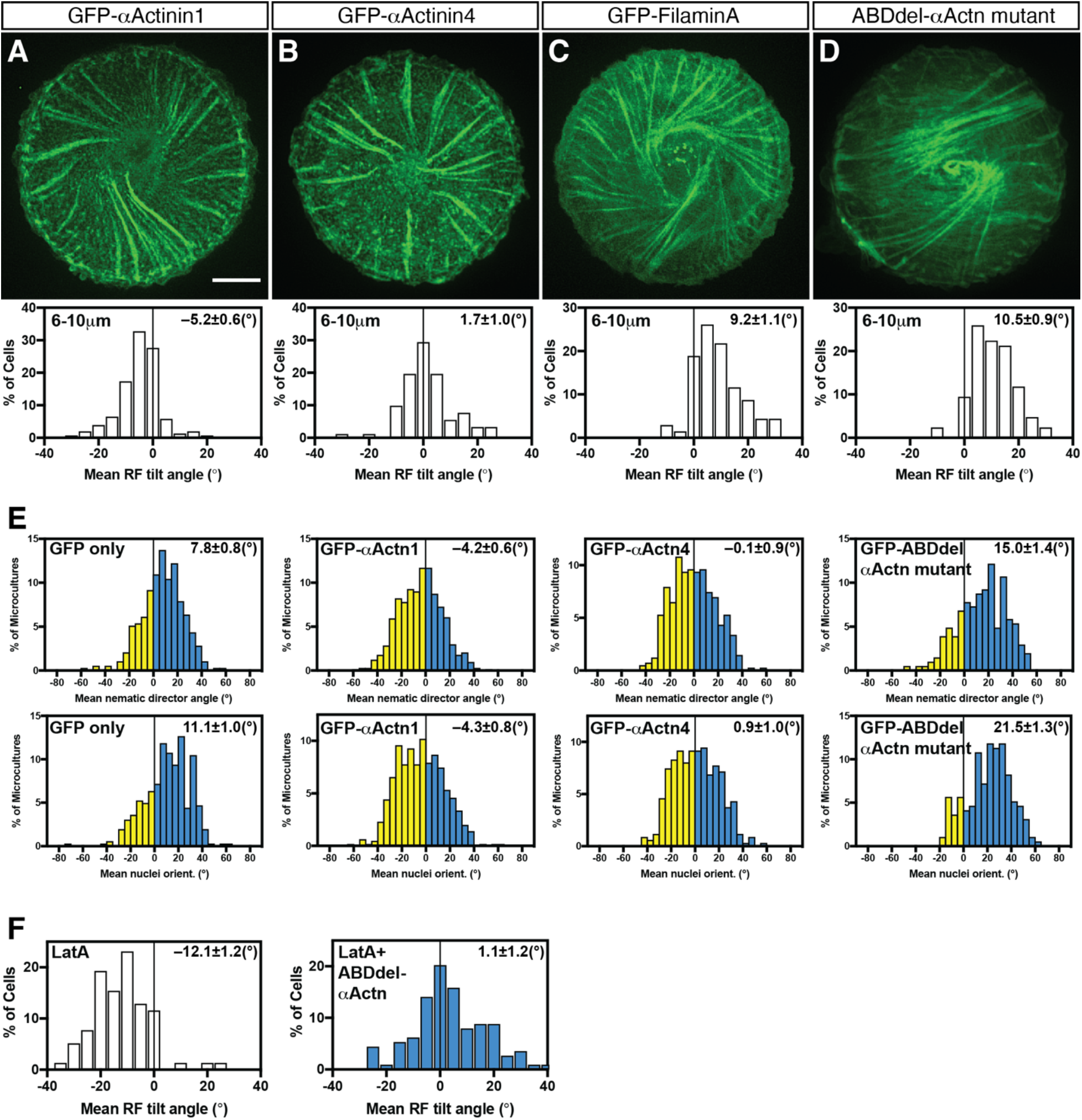
Effects of overexpression ofα-actinin-1 and other crosslinking proteins on radial fiber (RF) tilt and cell alignment in microcultures. Image of GFP-α-actinin-1 (**A**) expressing cell showing clockwise actin organization. Image of GFP-α-actinin-4 (**B**) expressing cell showing radial actin organization. Image of GFP-Filamin A (**C**) expressing cell showing anti-clockwise actin organization. (**D**) Anti-clockwise actin organization in GFP-ABDdel-α-actinin mutant expressing cell visualized by mRuby-LifeAct (pseudo-colored green). (**A** to **D**) The histograms show the distribution of average RF tilt in the 6-10 μm annulus in cells under corresponding conditions. See also Fig. 3, **B** and **F**. (**E**) Quantification of chiral alignment of cells transfected as indicated in microcultures as characterized by mean nematic directors (upper row) or mean nuclei orientation (lower row). The histograms were built based on average local cell orientation (nematic directors) values from 394 GFP-only transfected, 745 GFP-α-actinin-1 transfected, 417 GFP-α-actinin-4 transfected and 206 GFP-ABDdel-α-actinin mutant transfected microcultures respectively, or average nuclei orientation values from 364 GFP-only transfected cells, 659 GFP-α-actinin-1 transfected, 350 GFP-α-actinin-4 transfected and 195 GFP-ABDdel-α-actinin mutant transfected microcultures respectively. (**F**) Effect of inhibition of α-actinin-1 crosslinking function by GFP-ABDdel-α-actinin mutant on reversal of RF tilt by LatA treatment. Histograms showing the distribution of average RF tilt in the 6-10 μm annulus in 20nM LatA-treated cells (upper) (n= 78 cells) versus GFP-ABDdel-α-actinin mutant expressing cells treated with 20nM LatA (lower) (n= 114 cells). Mean±*s.e.m* values are indicated at the top right corner of each histogram. Scale bars, 10μm (**A** to **D**).

**Fig. S9.**
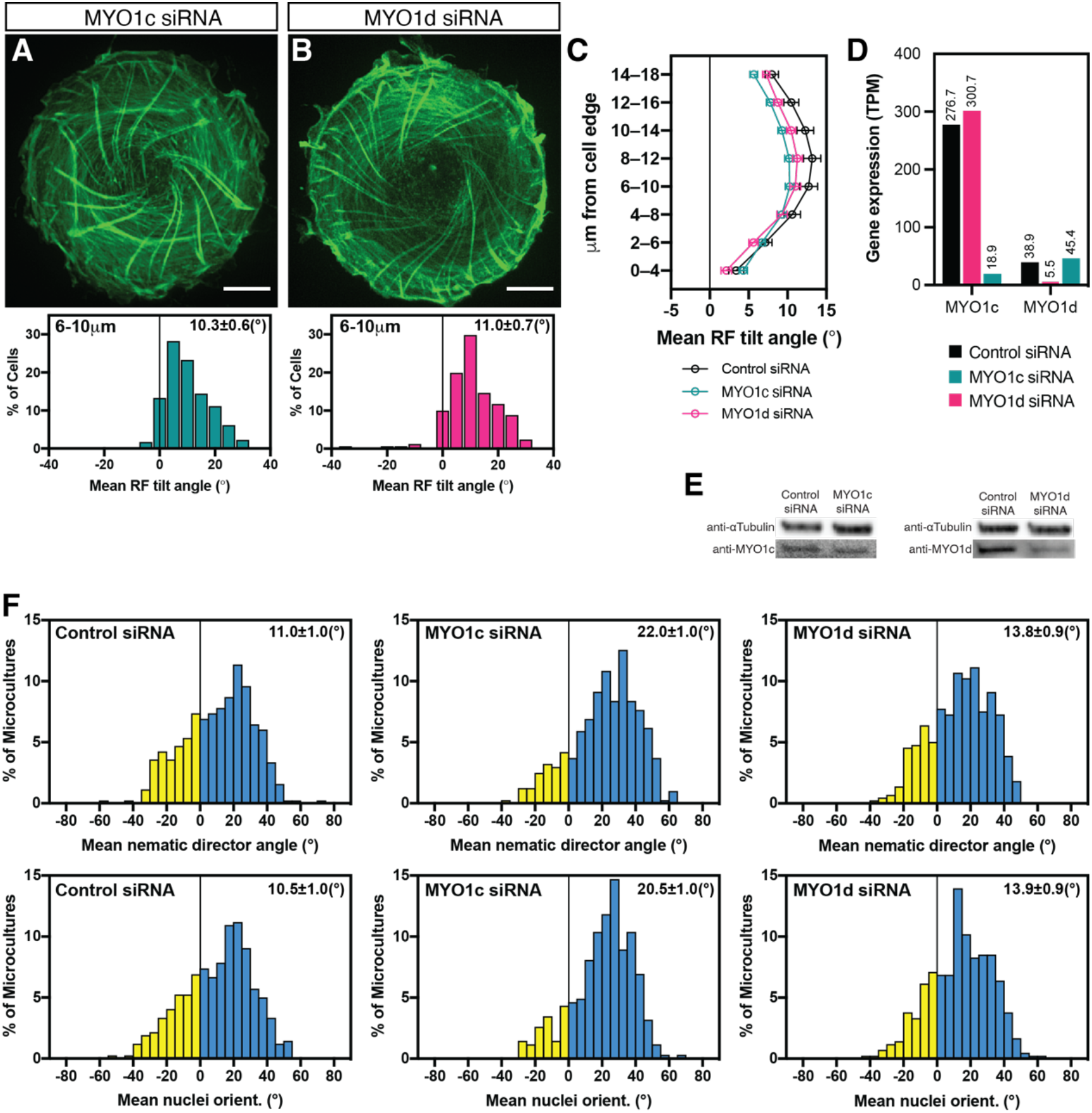
Left-right asymmetry of actin organization and chiral cell alignment in microcultures of myosin 1c and 1d knockdown cells. Actin organization visualized by phalloidin labeling in (**A**) myosin 1c (MYO1c) siRNA and (**B**) myosin 1d (MYO1d) siRNA transfected cells 6 hours following cell plating on circular pattern. The histograms show the distribution of average RF tilt in the 6-10 μm annulus in cells under corresponding conditions. (**C**) Average values of RF tilts (mean±*s.e.m*) as a function of the distance from the cell edge. (**D**) Transcriptome profiling of gene expression levels (transcripts per million; TPM) of MYO1c and MYO1d identified by RNA-sequencing in control-, MYO1c- and MYO1d- siRNA transfected human fibroblasts (n= 1 experiment). (**E**) Western blots showing MYO1c (left) or MYO1d (right) level in cells treated with scramble (control), anti-MYO1c or anti-MYO1d siRNA; α-tubulin was used as loading controls. (**F**) Quantification of chiral alignment of cells in microcultures as characterized by mean nematic directors (upper row) or mean nuclei orientation (lower row). Mean±*s.e.m* values are indicated at the top right corner of each histogram. Scale bars, 10μm (**A** and **B**). Sample sizes (n) for (**A**), (**B**), (**C**) and (**F**) can be found in Table S1.

**Fig. S10.**
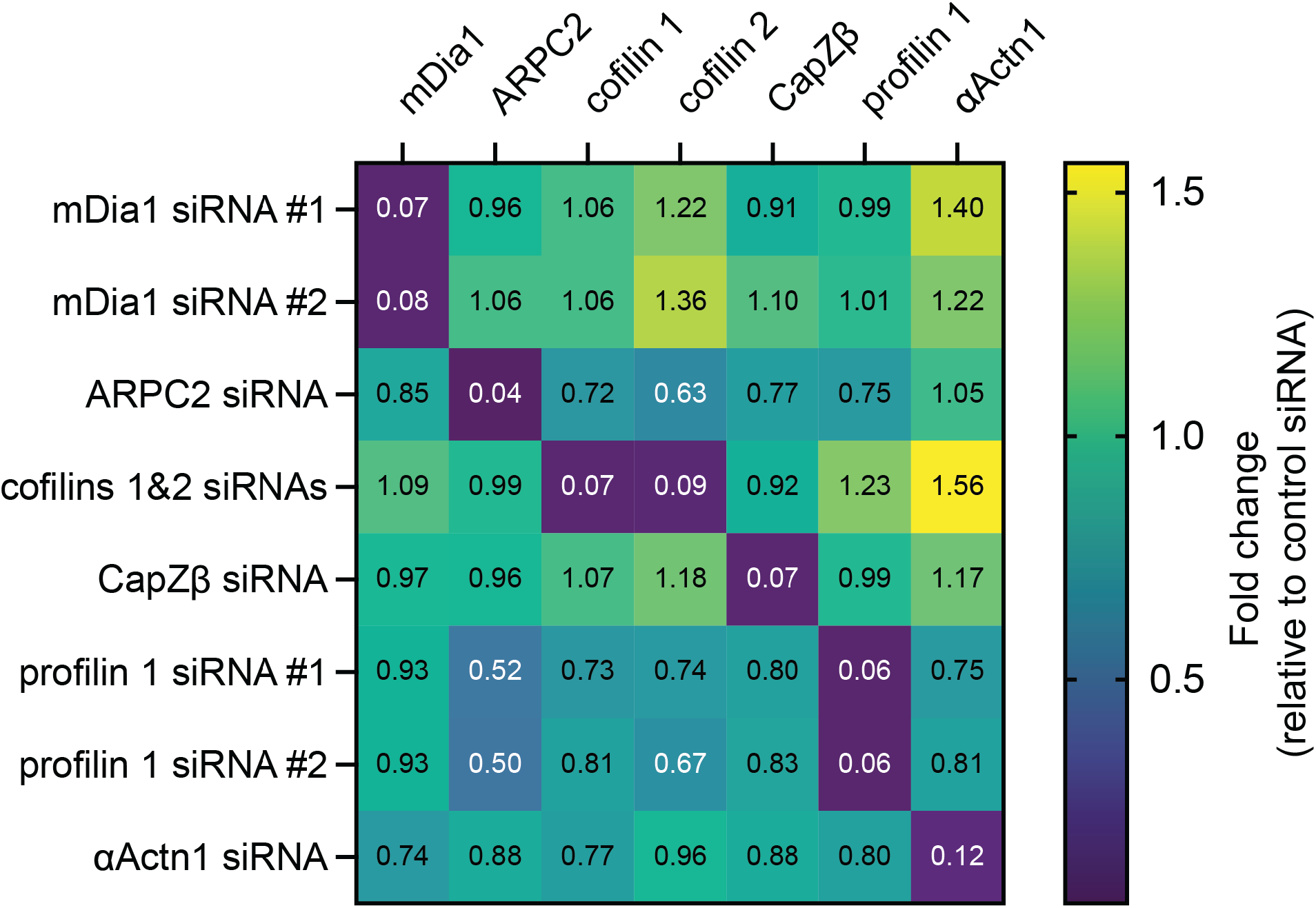
Transcriptional profile of major proteins associated with chirality phenotype of actin organization. Fold changes in gene expression levels (indicated by color coding and numbers) of major proteins in specific knockdown conditions as assessed by RNA-sequencing. #1 and #2 shows the results of two individual experiments with the same siRNA.

**Fig. S11.**
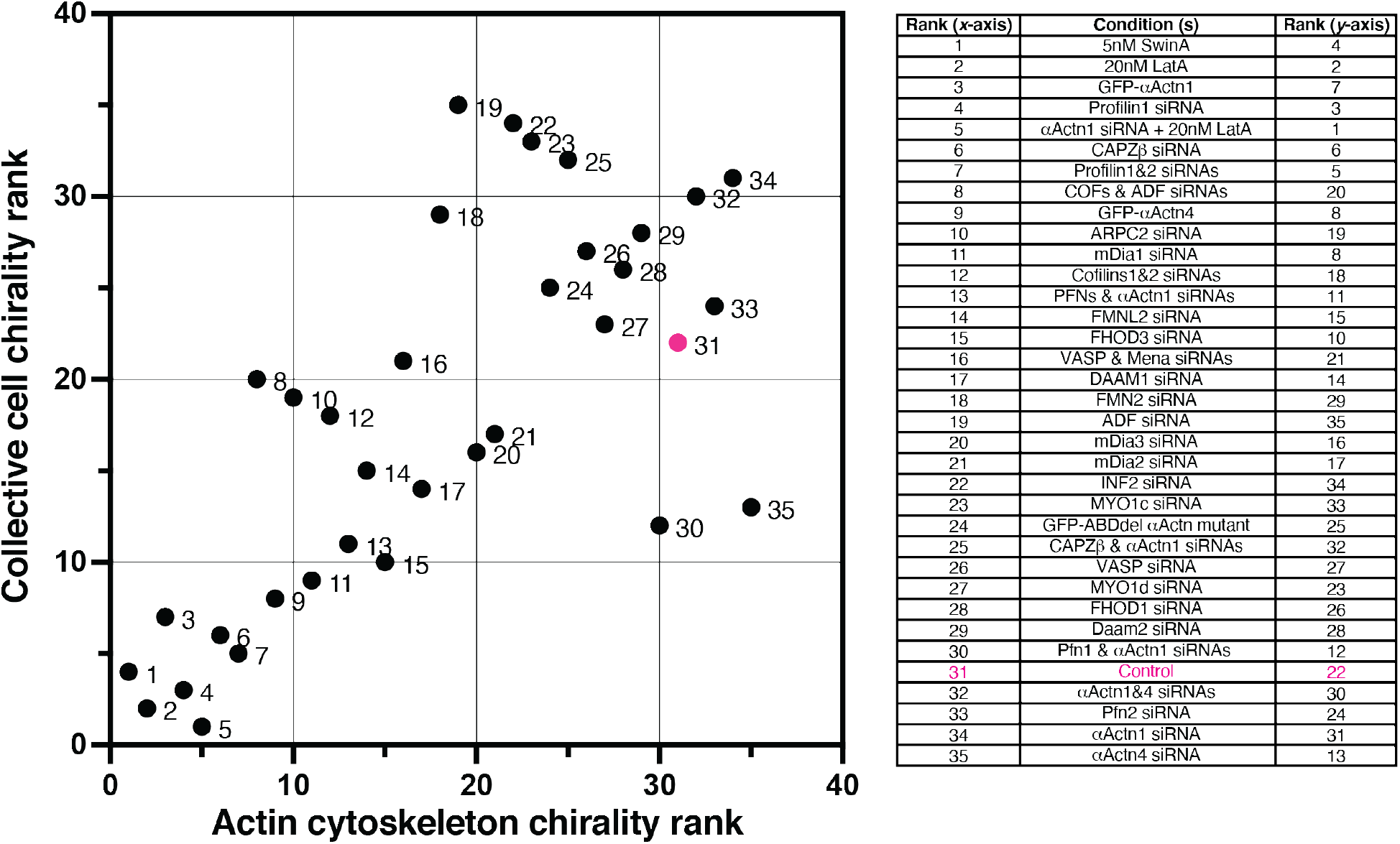
The rank correlation between actin cytoskeleton chirality in individual cells and collective cell chirality in microcultures. Each numbered dot represents average data from pooled experiments under respective conditions indicated in the list on the right. All dots are ranked according to actin cytoskeleton chirality value (*x*-axis) and collective chirality value (*y*-axis). Actin cytoskeleton chirality is defined as the mean radial fiber tilt angle at the 6-10 μm annulus. Collective chirality is defined as the mean nematic director angle for rectangular microcultures. Dots are indexed in ascending actin cytoskeleton chirality rank (*x*-axis). Position of control cell (31, 22) is shown in red. Numbers of cells and microcultures analyzed and the values of the means±*s.e.m* can be found in Table S1. See also Fig. 4.

## Notes

### Competing Interest Statement

The authors have declared no competing interest.

