## Supplementary Material for "Actin polymerization and crosslinking drive left-right asymmetry in single cell and cell collectives"

**This PDF file includes:**

Materials and Methods
Tables S1 to S3
Captions for Movies S1 to S6
References

**Other Supplementary Materials for this manuscript include the following:**

Movies S1 to S6

### Materials and Methods

Cells and plasmids transfection

Human foreskin fibroblasts (HFF) from American Type Culture Collection (catalog no. SCRC-1041) were cultured in Dulbecco's modified Eagle's medium high glucose supplemented with 10% fetal bovine serum (FBS), 1 mM sodium pyruvate and antibiotics (penicillin and streptomycin) at 5% $CO_2$ at 37°C. Enucleated cells were generated as described in our earlier work (*14*). Cells were transfected with DNA plasmids via electroporation (Neon® transfection system, Life Technologies) following manufacturer's instructions. Electroporation condition consists of two pulses of 1150 V for 30 milliseconds. Expression vectors encoding the following fluorescent fusion proteins were used: LifeAct-GFP (*11*), mRuby-LifeAct (*14*), mEmerald-mDia1-C-14 (Addgene plasmid # 54156), pEGFP-C1 (Clontech), GFP-α-actinin-1 (*11*), GFP-α-actinin-4 (gift of Dr. M. Pan, Mechanobiology Institute, Singapore), GFP-ABDdel-α-actinin-1 (*11*) (gift of Dr P. Roca-Cusachs, University of Barcelona, Barcelona, Spain), EGFP-Filamin A (gift of Dr. M. Sheetz, Mechanobiology Institute, Singapore), mCherry-Cofilin 1 (gift of Dr. C. G. Koh, Nanyang Technological University, Singapore) and mCherry-Profilin1 (Addgene plasmid #55121). Fluorescence-activated cell sorting (FACS) were performed to select for cells expressing high level of fluorescent fusion proteins of pEGFP-C1, GFP-α-actinin-1, GFP-α-actinin-4 or GFP-ABDdel-α-actinin-1. All cell culture and transfection reagents were obtained from Invitrogen. Other chemicals and reagents were obtained from Sigma, unless otherwise stated.

siRNA transfection

Cells were seeded into a 35mm dish on day 0 and transfected with 20 μM of siRNA using Lipofectamine RNAiMAX on days 1 and 2. For experiment involving individual cells, siRNA-transfected cells were trypsinized on day 4 and replated onto circular micropatterns. For experiment involving cell microcultures, siRNA-transfected cells were trypsinized on day 3 and replated onto rectangular micropatterns. As needed, transfection of plasmids via electroporation into siRNA-treated cells were performed on day 3 and cells were replated on day 4. siRNA transfected cells had their proteins or RNAs extracted on day 4 for immunoblotting or RNA sequencing respectively. siRNAs used in this study are listed in Table S2.

Micropatterning of substrates

Cells were seeded on substrates containing either circular micropattern of 1,800 $\mu m^2$ (individual cell experiment) or 300×600 μm rectangles (multicellular microculture experiment). Each micropatterned substrate was fabricated by stencil patterning as previously described in our earlier work (*14*). Briefly, a PDMS stamp was generated as described in (*11*). The PDMS stamp was then inverted and placed onto a hydrophobic uncoated 35 mm μ-dish (ibidi GmbH). Norland Optical Adhesive 73 (NOA-73, Norland Inc.) was deposited along an edge of the stamp and allowed to flow through the gaps between the PDMS stamp and dish by capillary action, upon which the stamp was sealed on all sides using NOA-73. The NOA-73 stencil was cured under ultraviolet illumination for 15 s. After peeling off the PDMS stamp, the stencil and dish were incubated with fibronectin (Calbiochem, Merck Millipore) at a concentration of 50 $\mu g\ ml^{-1}$ in 1×PBS at 4°C overnight after a brief degassing at 10 mbar. At the end of the incubation, the fibronectin solution was aspirated, and the stencil was removed. The printed dish bottom was passivated with 0.2% Pluronic acid-$H_2O$ for 10 min. Finally, the passivated dishes were washed thrice with 1×PBS before cell seeding.

Assessment of individual cells on circular micropattern

Cells were seeded on printed dishes containing circular micropatterns at a density of $5\times10^4$ cells ml$^{-1}$ for 10 min. The medium containing unattached cells was then replaced with fresh DMEM. After 6 hours' incubation, the cells were fixed using 4% paraformaldehyde (Tousimis, USA) in PBS for 15 minutes, followed by three 1×PBS washes. Cells were permeabilised using 0.1% Triton-X-100 in PBS, and then blocked with 2% bovine serum albumin (BSA)-PBS for 1 h at room temperature before incubation with appropriate labelling reagents. Actin and nucleus staining were performed using phalloidin (Molecular Probes) and Hoechst 33342 (Invitrogen) respectively. For live cell imaging experiment, cells were seeded on circular micropatterns at a density of $5\times10^4$ cells ml$^{-1}$ for 10 min. The medium containing unattached cells was then replaced with Leibovitz's L-15 containing 10% FBS. Cells were left for at least 2 h before imaging at 37°C with 5% $CO_2$. Time-lapse images at 10-20 min intervals and Z-stacks of step-size 0.35 µm with total height of 10-15 µm were acquired with a spinning disc confocal microscope (PerkinElmer Ultraview VoX) attached to an Olympus IX81 inverted microscope, equipped with a 100× oil immersion objective (1.40 NA, UPlanSApo), an EMCCD camera (C9100-13, Hamamatsu Photonics) for image acquisition, and Volocity software (PerkinElmer) to control the set-up. Fixed samples were also imaged with the same step-up. Maximum projection of the Z-stack images was performed with Volocity software or with Fiji software and exported as 16-bit TIFF files (512×512 pixel and 0.138502 µm pixel$^{-1}$). Each image contained a single cell and these images were subsequently used for deep learning-based identification of radial fibers.

Assessment of multicellular microcultures on rectangular micropatterns

Cells were seeded on rectangular micropatterns at a density of $1\times10^5$ cells ml$^{-1}$ for 20 min. The medium containing unattached cells was then replaced with fresh DMEM and cell microcultures were incubated for a total of 48 hours before cell fixation using 4% paraformaldehyde-PBS for 15 minutes. Just prior to fixation, cell nuclei were stained with 1 µg ml$^{-1}$ Hoechst 33342 for 10 min. Brightfield and widefield images of cell microcultures were taken using a 20× air objective (0.45 NA, LUCPLFLN20X, Olympus) on an Olympus IX81 inverted microscope, equipped with Andor Neo 5.5 sCMOS camera and light source (Lumencor SOLA SE Light Engine). Single plane images of phase contrast and DAPI channels were taken. Each image contained a single rectangular cell microculture and these images were subsequently used for measurement of average nematic directors and nuclei orientation angles in rectangular microcultures.

Drug treatment

For drug inhibition studies, following cell seeding on micropatterns, the medium containing unattached cells was then replaced with fresh medium containing either 20 nM latrunculin A (Santa Cruz Biotechnology Inc., SCB Inc.) or 5 nM swinholide A (SCB Inc.). For experiments that lasted more than 24 h, fresh drugs were added every 24 h until the end of the observation period. All inhibitors remained in the medium during the entire period of observation, except in drug washout experiments.

Segmentation of radial fibers

Images of the actin cytoskeleton were first converted to 8-bit and the 'Enhance brightness/contrast' function in Fiji was used with the 'saturated pixels' parameter set to the default of 0.35. A Unet-ResNet50 deep learning model (*41*), implemented in Python, was trained to identify actin radial fibers in cells confined on circular micropattern. Briefly, the following steps were taken. The model was trained using 32 images of actin cytoskeleton labelled by fluorescent protein tagged-LifeAct. These training images comprise of cells with their actin cytoskeleton in a radial or chiral organization, and images of varied intensities were selected. Data augmentation was done using the Albumentations library (*42*). The ground-truths (binary, 8-bit) were prepared by manual demarcation of actin radial fibers in Fiji. The code and complete list of parameters of the trained deep learning model is available upon request. This trained model was used to identify radial fibers in both phalloidin- and LifeAct-labelled cells, returning a 32-bit image of identified radial fibers. Segmentation of these identified radial fibers was performed using a custom MATLAB script, in which background subtraction (with rolling ball of 30-pixel radius) and then Niblack local thresholding (*43*), with window size of 15×15, k = -0.3 and offset = -0.01, were applied. The resulting binary image was then skeletonized using the MATLAB built-in function, bwmorph. Intersecting radial fibers were separated by branch point removal, and fiber segments with similar orientation (angle difference ≤ 30º) and at nearby position (distance ≤ 30 pixels) were connected as a single fiber.

Measurement of radial fiber tilt angles

The following procedures were performed using a custom MATLAB script unless otherwise stated. Cell segmentation was performed by thresholding the Gaussian-smoothed (sigma value set to 3) actin image using Otsu binarization (threshold value set to 0.4 of Otsu auto threshold), followed by a series of mathematical morphological operations (imclose, imfill, imerode). Cell centroid and cell spread area were calculated based on this cell mask. The cell mask was also used for generating concentric ring masks of 4 μm in width starting from the cell edge, with 2 μm increments, for masking the segmented radial fibers. In each ring, the angle of each radial fiber segment was measured relative to the cell edge. The angle at the cell edge θ, computed using Python, was given by the formula $\theta = \arcsin\left(\frac{r \sin\, \theta_r}{R}\right)$, where $\vec{R}$ connects the cell centroid and intersection of the continuation of the radial fiber segment with the edge of the cell and $\vec{r}$ connects the cell centroid and intersection of radial fiber with outer edge of the annulus. $\theta$ and $\theta_r$ are the angles between the radial fiber segment and $\vec{R}$ and $\vec{r}$ respectively. See also Fig. S1. Based on visual inspection, actin radial fiber segments with $\theta_r$ more than or equals to 68º were unlikely to be radial fibers and were omitted from the analysis. The average inflation of area of the cell mask relative to the area of the micropatterns (1800 μm$^2$) was estimated to be 63.353 μm$^2$ using a dataset of ~100 cells. This constant was subtracted from the cell area before the computation of cell radius R. Only cells with area between 1700 and 2000 μm$^2$ were analyzed.

Measurement of average nematic directors and nuclei orientation angles in rectangular microcultures

The following procedures were performed using a custom MATLAB script unless otherwise stated. First, identification and segmentation of individual rectangular microculture using phase contrast images was done by performing a Wiener filter with a neighborhood size of 20×20 pixels to remove image noise. This was followed by an entropy filter with a 3×3 pixel structural element and morphological opening with a 9×9 pixel structural element. This results in an image that differentiates between areas with and without cells. Otsu binarization was then performed to segment the image, the segmented area at the center of the image was selected as the segmentation mask. This serves as an indicator of the area covered with cells. The bounding box enclosing this segmented area serves to represent the dimensions of the microculture. Only microcultures with bounding box width of 225 to 375 µm and height of more than 550 µm, and with a segmentation mask that covered more than 80% of the bounding box area were analyzed. For each bounding box, the center 200×500 µm region of interest was used for subsequent steps in the measurements of average nuclei orientation and average nematic director orientation.

Segmentation of the nuclei was achieved using NICK adaptive binarization (*44, 45*). A Wiener filter using a neighborhood size of 9×9 pixels was performed prior to segmentation. The concaved-point based splitting algorithm, described by (*46*), was used to separate any overlapping nuclei. Segmented objects above the size of 10000 pixels were removed as these corresponded to the background regions, while objects smaller than 500 pixels were also removed as these were either fragmented nuclei or noise regions that were segmented by chance. In addition, only nucleus that had a centroid position that laid within the bounding box of the segmented phase contrast image was selected for further analysis. The number of nuclei in the bounding box was also counted. Microcultures that had less than 50 nuclei were removed as these microcultures often had too few cells to cover the entire rectangular micropattern. The orientation of these nuclei was then calculated based on the angle of the long axis of a fitted ellipse with respect to the long axis of the rectangular micropattern. Alignment of the cell group was determined based on the mean resultant length (*47*) of the nuclei orientation. A cutoff value of 0.35 was selected and any rectangle with a mean resultant length greater than that was classified as aligned. The mean nuclei orientation per aligned microculture was determined by calculating the mean of all the orientations of the individual nuclei within a single microculture.

Local cell orientation in the phase contrast image was calculated by obtaining the nematic director field as described in (*26*). Briefly the orientation tensor was obtained using OrientationJ implemented in Fiji and the nematic director was obtained using a window size of 60×60$\mu m^2$ and 70% overlap. The orientation of each directors was measured as the angle relative to the long axis of the rectangular micropattern. The orientation of the directors within the region of interest was then used to determine the alignment of the microculture in a similar manner as that for the nuclei orientation. A higher cutoff value of 0.5 for alignment was set due to more coherent nature of the nematic directors. The mean nematic director angle per aligned microculture was determined by calculating the mean of all the orientations of the directors within a single microculture.

<u>Immunoblotting</u>

Cell pellets collected in RIPA buffer (SCB Inc.), containing 2 µL $ml^{-1}$ protease inhibitors cocktail (Sigma, catalogue no. P8340), and were mechanically lysed by syringing through a 27.25G needle on ice. Protein concentration was quantified using the Micro BCA Protein Assay Kit (Thermo Scientific) according to manufacturer's instructions. 20 µg of protein lysate was dissolved in 1×Laemmli sample buffer supplemented with 5% 2-mercaptoethanol, and separated by 4-20% SDS-polyacrylamide gel (GenScript USA Inc) electrophoresis at 100V for 1 hour and then transferred to a 0.4 µm pore size PVDF membrane (Thermo Scientific, catalog number 88518) at 100V for 2 hours for formin proteins and 1 hour for other proteins in an ice bath. The PVDF membrane was blocked using Intercept® (TBS) Blocking Buffer (LI-COR, Inc.) for 1 hour at room temperature before incubation at 4 ºC overnight with appropriate primary antibodies. Primary antibodies were diluted in blocking buffer containing 0.1% Tween-20 at their respective concentrations summarized in Table S3. After washes in TBS-T, the membrane was probed with either IRDye® 680RD Goat anti-Rabbit IgG (LI-COR, dilution 1:10 000) or IRDye® 800CW Goat anti-Mouse IgG (LI-COR, dilution 1:15 000) for 1 hour at room temperature. The membrane was then washed in TBS-T before fluorescent detection with an Odyssey® CLx imaging system at a resolution of 169 µm and 'medium' quality settings on Image Studio software.

Transcriptome profiling by RNA sequencing

RNAs were extracted using RNeasy® Plus Universal Kits (Qiagen) according to manufacturer's instructions. Library is prepared using TruSeq Stranded mRNA LT Sample Prep Kit and sequenced using NovaSeq6000 Illumina platform. Alignment was performed (STAR aligner) and trimmed reads were mapped to GRCh38 reference genome with HISAT2, splice-aware aligner. Gene expression was expressed using transcript per million reads.

Image and statistical analysis

Image processing and analysis were performed with Fiji software, MATLAB and Python. All customized image analysis scripts are available upon request. Numbers of samples (n) measured are specified in figure legends for all of the quantitative data and in Table S1. Prism (GraphPad Software) was used for statistical analysis. No statistical method was used to predetermine sample size.

**Table S1. Mean radial fiber (RF) tilt angles at the 6-10 μm annulus and corresponding mean nematic director angles for rectangular microcultures for each type of treatment.** Ranks and sample sizes (n) are included.

| Rank of RF tile angle | Condition(s) | Mean RF tilt angle (°) | Mean nematic director angle (°) | Rank of nematic director angle | n= (cells) | n= (micro-cultures) |
|---|---|---|---|---|---|---|
| 1 | 5nM SwinA | -11.80±0.78 | -13.43±0.70 | 4 | 153 | 1108 |
| 2 | 20nM LatA | -10.24±0.48 | -22.47±0.54 | 2 | 490 | 1416 |
| 3 | GFP-αActn1 | -5.18±0.60 | -4.15±0.65 | 7 | 156 | 745 |
| 4 | Profilin1 siRNA | -4.59±0.51 | -17.50±0.77 | 3 | 431 | 715 |
| 5 | αActn1 + 20nM LatA | -2.38±1.55 | -26.24±1.20 | 1 | 94 | 237 |
| 6 | CAPZβ siRNA | -2.21±0.52 | -7.56±0.72 | 6 | 409 | 661 |
| 7 | Profilins1&2 siRNAs | -0.37±1.08 | -8.59±1.38 | 5 | 76 | 207 |
| 8 | COFs & ADF siRNAs | -0.11±0.41 | 11.16±1.19 | 20 | 308 | 216 |
| 9 | GFP-αActn4 | 1.68±0.97 | -0.07±0.88 | 8 | 92 | 417 |
| 10 | ARPC2 siRNA | 2.59±0.53 | 10.38±1.05 | 19 | 434 | 342 |
| 11 | mDia1 siRNA | 2.78±0.36 | 4.02±0.71 | 9 | 515 | 894 |
| 12 | Cofilins1&2 siRNA | 3.37±0.43 | 10.00±0.99 | 18 | 426 | 373 |
| 13 | PFNs & αActn1 siRNAs | 4.45±0.76 | 6.39±1.34 | 11 | 295 | 227 |
| 14 | FMNL2 siRNA | 6.66±0.64 | 8.87±0.95 | 15 | 227 | 454 |
| 15 | FHOD3 siRNA | 7.08±0.64 | 5.99±0.87 | 10 | 262 | 407 |
| 16 | VASP & Mena siRNAs | 7.63±0.76 | 12.26±1.10 | 21 | 143 | 366 |
| 17 | DAAM1 siRNA | 7.78±0.70 | 8.21±1.44 | 14 | 268 | 165 |
| 18 | FMN2 siRNA | 7.97±0.92 | 16.02±1.49 | 29 | 115 | 181 |
| 19 | ADF siRNA | 9.38±0.54 | 24.36±1.01 | 35 | 339 | 295 |
| 20 | mDia3 siRNA | 9.58±0.99 | 9.05±2.93 | 16 | 98 | 32 |
| 21 | mDia2 siRNA | 9.93±0.60 | 9.41±1.04 | 17 | 283 | 419 |
| 22 | INF2 siRNA | 9.95±0.53 | 22.56±0.74 | 34 | 356 | 495 |
| 23 | MYO1c siRNA | 10.29±0.58 | 21.99±0.98 | 33 | 181 | 406 |
| 24 | GFP-ABDdel αActn mutant | 10.51±0.87 | 14.99±1.37 | 25 | 85 | 206 |
| 25 | CAPZβ & αActn1 siRNAs | 10.69±0.73 | 18.73±0.78 | 32 | 219 | 360 |
| 26 | VASP siRNA | 10.84±0.46 | 15.70±1.78 | 27 | 395 | 202 |
| 27 | MYO1d siRNA | 11.01±0.69 | 13.78±0.86 | 23 | 171 | 440 |
| 28 | FHOD1 siRNA | 11.45±0.91 | 15.03±1.26 | 26 | 147 | 157 |

| | | | | | | |
|---|---|---|---|---|---|---|
| 29 | DAAM2 siRNA | 11.66±1.10 | 15.92±1.25 | 28 | 100 | 215 |
| 30 | Pfn1 & αActn1 siRNAs | 11.84±1.33 | 7.90±1.67 | 12 | 42 | 135 |
| 31 | Control | 11.85±0.25 | 12.56±0.28 | 22 | 1559 | 4432 |
| 32 | αActn1&4 siRNAs | 15.47±1.22 | 16.19±1.07 | 30 | 105 | 182 |
| 33 | Profilin2 siRNA | 17.16±0.63 | 14.98±0.76 | 24 | 182 | 519 |
| 34 | αActn1 siRNA | 17.35±0.62 | 17.61±0.78 | 31 | 264 | 430 |
| 35 | αActn4 siRNA | 18.00±0.70 | 8.20±1.47 | 13 | 116 | 135 |

**Table S2. List of siRNAs used.**

| siRNA | Company, Product name or Target Sequence(s) | Catalog No. |
|---|---|---|
| Control | Dharmacon, ON-TARGETplus Non-targeting control | D-001810-01 |
| Alpha-Actinin1 | Dharmacon, ON-TARGETplus SMARTpool, Human ACTN1 siRNA | L-011195-00 |
| Alpha-Actinin4 | Dharmacon, ON-TARGETplus SMARTpool, Human ACTN4 siRNA | L-011988-00 |
| ADF | Dharmacon, ON-TARGETplus, Human DSTN siRNA | J-012303-05 & J-012303-06 |
| ARPC2 | Dharmacon, ON-TARGETplus SMARTpool, Human ARPC2 siRNA | L-012081-00 |
| CapZβ | Dharmacon, ON-TARGETplus SMARTpool, Human CAPZB siRNA | L-011990-00 |
| Cofilin 1 | Santa Cruz Biotechnology Inc, Cofilin 1 siRNA (h) | sc-35078 |
| Cofilin 2 | Santa Cruz Biotechnology Inc, Cofilin 2 siRNA (h) | sc-37027 |
| DAAM1 | Santa Cruz Biotechnology Inc, DAAM1 siRNA (h) | sc-62190 |
| DAAM2 | Santa Cruz Biotechnology Inc, DAAM2 siRNA (h) | sc-62192 |
| FHOD1 | Santa Cruz Biotechnology Inc, FHOD1 siRNA (h) | sc-60635 |
| FHOD3 | Dharmacon, ON-TARGETplus SMARTpool, Human FHOD3 siRNA | L-023411-01 |
| FMNL2 | Dharmacon, ON-TARGETplus, Human FMNL2 siRNA | J-031993-09 & J-031993-10 |
| FMN2 | Santa Cruz Biotechnology Inc, Formin 2 siRNA (h) | sc-43765 |
| INF2 | Santa Cruz Biotechnology Inc, INF2 siRNAs (h) | sc-92159 |
| mDia1 | Dharmacon, ON-TARGETplus, Human DIAPH1 siRNA | J-010347-06 |
| mDia2 | Dharmacon, ON-TARGETplus SMARTpool, Human DIAPH3 siRNA | L-018997-00 |
| mDia3 | Dharmacon, ON-TARGETplus, Human DIAPH2 siRNA | J-012029-05 & J-012029-06 |
| Mena | Santa Cruz Biotechnology Inc, Mena siRNA (h) | sc-43496 |
| MYO1c | Santa Cruz Biotechnology Inc, Myosin 1c siRNA (h) | sc-44604 |
| MYO1d | Santa Cruz Biotechnology Inc, Myosin 1d siRNA (h) | sc-44608 |
| Profilin 1 | 5’-GCAAAGACCGGUCAAGUUU-3’ and 5’-CACGGUGGUUUGAUCAACA-3’ | |
| Profilin 2 | 5’-GUAGAGCAUUGGUUAUAGU-3’ and 5’-CCAGGGACAUUCCAUCAUU-3’ | |
| VASP | Santa Cruz Biotechnology Inc, VASP siRNA (h) | sc-29516 |

**Table S3. List of antibodies used.**

| Protein | Company | Catalog No. | Dilution |
|---|---|---|---|
| α-Tubulin | Sigma-Aldrich | T5168 | 1:5000 |
| ADF | Abcam | ab186754 | 1:1000 |
| ARPC2 | Santa Cruz Biotechnology Inc | sc-515754 (F-5) | 1:1000 |
| CAPZβ | Abcam | ab175212 | 1:1000 |
| Cofilins1&2 | Santa Cruz Biotechnology Inc | sc-376476 (E-8) | 1:1000 |
| DAAM1 | Abcam | ab56951 | 1:1000 |
| DAAM2 | Santa Cruz Biotechnology Inc | sc-515129 (E-1) | 1:1000 |
| FHOD1 | ECM Biosciences | FM3521 | 1:1000 |
| FMN2 | Santa Cruz Biotechnology Inc | sc-376787 (C-3) | 1:1000 |
| GAPDH | Santa Cruz Biotechnology Inc | sc-47724 (0411) | 1:5000 |
| INF2 | Proteintech | 20466-1-AP | 1:1000 |
| mDia1 | BD Biosciences | 610849 | 1:1000 |
| mDia3 | ECM Biosciences | DP4511 | 1:1000 |
| Mena | Santa Cruz Biotechnology Inc | sc135988 (21) | 1:1000 |
| MYO1c | Santa Cruz Biotechnology Inc | sc-136544 (13) | 1:1000 |
| MYO1d | Santa Cruz Biotechnology Inc | sc-515292 (H-1) | 1:1000 |
| ARPC2 | Santa Cruz Biotechnology Inc | sc-515754 (F-5) | 1:1000 |
| Profilin 1 | Santa Cruz Biotechnology Inc | sc-137235 (B-10) | 1:1000 |
| Profilin 2 | Santa Cruz Biotechnology Inc | sc-100955 (4K-6) | 1:1000 |
| VASP | Santa Cruz Biotechnology Inc | sc-46668 (A-11) | 1:1000 |

### Movies Captions

**Movie S1.** Anti-clockwise (dextral) chiral actin swirling in control HFF cell confined to circular fibronectin pattern. LifeAct labeling of actin (yellow) and Hoechst 33342 labeling of the nucleus (magenta) are shown. Images were recorded at 20 minute intervals over a period of 14 hours. Display rate is 7 frames/sec.

**Movie S2.** Left-right asymmetric alignment of control HFF cells plated on rectangular fibronectin pattern (300×600 μm). Phase-contrast microscopy. Images were recorded at 2 hours intervals over a period of 56 hours. Display rate is 3 frames/sec.

**Movie S3.** Clockwise (sinistral) chiral actin swirling in profilin 1 siRNA knockdown HFF cell confined to circular fibronectin pattern. LifeAct labeling of actin (yellow) is shown. Images were recorded at 30 minute intervals over a period of 14 hours. Display rate is 7 frames/sec and corresponds to the cell shown in Fig. 2B.

**Movie S4.** Reversal of swirling direction upon addition of 20nM of latrunculin A (LatA). Time of addition of drug is indicated. LifeAct labeling of actin (yellow) is shown. Images were recorded at 2 minute intervals over a period of ~ 2 hours. Display rate is 7 frames/sec.

**Movie S5.** Reversal of swirling direction upon latrunculin A washout. Time of removal of drug is indicated. LifeAct labeling of actin (yellow). Images were recorded at 3 minute intervals over a period of ~ 4 hours. Display rate is 7 frames/sec.

**Movie S6.** Anti-clockwise actin swirling in enucleated HFF cell (left) and clockwise actin swirling in enucleated HFF cell in the presence of 20nM latrunculin A (right). LifeAct labeling of actin (green). Images were recorded at 10 minute intervals over a period of 9.5 hours. Display rate is 7 frames/sec and corresponds to the cells shown in Fig. S7, B and D.
